# Unsupervised clustering of temporal patterns in high-dimensional neuronal ensembles using a novel dissimilarity measure

**DOI:** 10.1101/252791

**Authors:** Lukas Grossberger, Francesco P. Battaglia, Martin Vinck

**Affiliations:** Donders Institute for Brain, Cognition and Behaviour, Radboud Universiteit, Nijmegen, The Netherlands; Ernst Strüngmann Institute for Neuroscience in Cooperation with Max Planck Society, Frankfurt am Main, Germany

## Abstract

Temporally ordered multi-neuron patterns likely encode information in the brain. We introduce an unsupervised method, SPOTDisClust (Spike Pattern Optimal Transport Dissimilarity Clustering), for their detection from high-dimensional neural ensembles. SPOTDisClust measures similarity between two ensemble spike patterns by determining the minimum transport cost of transforming their corresponding normalized cross-correlation matrices into each other (SPOTDis). Then, it performs density-based clustering based on the resulting inter-pattern dissimilarity matrix. SPOTDisClust does not require binning and can detect complex patterns (beyond sequential activation) even when high levels of out-of-pattern “noise” spiking are present. Our method handles efficiently the additional information from increasingly large neuronal ensembles and can detect a number of patterns that far exceeds the number of recorded neurons. In an application to neural ensemble data from macaque monkey V1 cortex, SPOTDisClust can identify different moving stimulus directions on the sole basis of temporal spiking patterns.

**Author summary:** The brain encodes information by ensembles of neurons, and recent technological developments allow researchers to simultaneously record from over thousands of neurons. Neurons exhibit spontaneous activity patterns, which are constrained by experience and development, limiting the portion of state space that is effectively visited. Patterns of spontaneous activity may contribute to shaping the synaptic connectivity matrix and contribute to memory consolidation, and synaptic plasticity formation depends crucially on the temporal spiking order among neurons. Hence, the unsupervised detection of spike sequences is a sine qua non for understanding how spontaneous activity contributes to memory formation. Yet, sequence detection presents major methodological challenges like the sparsity and stochasticity of neuronal output, and its high dimensionality. We propose a dissimilarity measure between neuronal patterns based on optimal transport theory, determining their similarity from the pairwise cross-correlation matrix, which can be taken as a proxy of the “trace” that is left on the synaptic matrix. We then perform unsupervised clustering and visualization of patterns using density clustering on the dissimilarity matrix and low-dimensional embedding techniques. This method does not require binning of spike times, is robust to noise, jitter and rate fluctuations, and can detect more patterns than the number of neurons.

## 1 Introduction

Precisely timed spike patterns spanning multiple neurons are a ubiquitous feature of both spontaneous and stimulus-evoked brain network activity. Remarkably, not all patterns are generated with equal probability. Synaptic connectivity, shaped by development and experience, favors certain spike sequences over others, limiting the portion of the network’s “state space” that is effectively visited (Luczak, McNaughton, and Harris, 2015; Ikegaya et al., 2004). The structure of this permissible state space is of the greatest interest for our understanding of neural network function. Multi-neuron temporal sequences encode information about stimulus variables (Vinck et al., 2010; Siegel, Warden, and Miller, 2009; Havenith et al., 2011; Konig et al., 1995; Lu, Liang, and Wang, 2001; Gerstner et al., 1996), in some cases “unrolling” non-temporally organized stimuli, such as odors, into temporal sequences (Wehr and Laurent, 1996). Recurrent neuronal networks can generate precise temporal sequences (Memmesheimer et al., 2014; Abbott and Blum, 1996; Huh and Sejnowski, 2017; Fiete et al., 2010; Laje and Buonomano, 2013), which are required for example for the generation of complex vocalization patterns like bird songs (Hahnloser, Kozhevnikov, and Fee, 2002). Temporal spiking patterns may also encode sequences of occurrences or actions, as they take place, or are planned, projected, or “replayed” for memory consolidation in the hippocampus and other structures (Carr, Jadhav, and Frank, 2011; Foster and Wilson, 2006; Dragoi and Tonegawa, 2011; Johnson and Redish, 2007; Nadasdy et al., 1999; Pfeiffer and Foster, 2013; Skaggs and McNaughton, 1996; Euston, Tatsuno, and McNaughton, 2007; Peyrache et al., 2009; Pastalkova et al., 2008).

Timing information between spikes of different neurons is critical for memory function, as it regulates spike timing dependent plasticity (STDP) of synapses, with firing of a post-synaptic neuron following the firing of a pre-synaptic neuron typically inducing synaptic potentiation, and firing in the reverse order typically inducing depotentiation (Dan and Poo, 2004; Markram et al., 1997; Abbott and Nelson, 2000). Thus, the consolidation of memories may rely on recurring temporal patterns of neural activity, which stabilize and modify the synaptic connections among neurons (Buzsaki, 1989; Carr, Jadhav, and Frank, 2011; Foster and Wilson, 2006; Dragoi and Tonegawa, 2011; Johnson and Redish, 2007; Nadasdy et al., 1999; Pfeiffer and Foster, 2013; Skaggs and McNaughton, 1996; Benchenane et al., 2010; Sejnowski and Paulsen, 2006; Suri and Sejnowski, 2002; Lee and Wilson, 2002; Drew and Abbott, 2006; van Rossum, Bi, and Turrigiano, 2000). Storing memories as sequences has the advantage that a very large number of patterns is possible, because the number of possible spike orderings grows exponentially, and different sequences can efficiently be associated to different memory items, as proposed by for instance the reservoir computing theory (Maass, Natschlager, and Markram, 2002; Lazar, Pipa, and Triesch, 2009; Singer and Lazar, 2016; Buonomano and Maass, 2009; Lukoševičius and Jaeger, 2009).

Detecting these temporal patterns represents a major methodological challenge. With recent advances in neuro-technology, it is now possible to record from thousands of neurons simultaneously (Jun et al., 2017), and this number is expected to show an exponential growth in the coming years (Stevenson and Kording, 2011). The high dimensionality of population activity, combined with the sparsity and stochasticity of neuronal output, as well as the limited amount of time one can record from a given neuron, makes the detection of recurring temporal sequences an extremely difficult computational problem. Many approaches to this problem are supervised, that is, they take patterns occurring concurrently with a known event, such as the delivery of a stimulus for sensory neurons or the traversal of a running track for hippocampal place fields, as a “template” and then search for repetitions of the same template in spiking activity (Lee and Wilson, 2004; Nadasdy et al., 1999; Davidson, Kloosterman, and Wilson, 2009). Other approaches construct a template by measuring latencies of each neuron’s spiking from a known event, such as the beginning of a cortical UP state (Havenith et al., 2011; Luczak, Bartho, and Harris, 2009). While this enables rigorous, relatively easy statistical treatment, it risks neglecting much of the structure in the spiking data, which may contain representations of other items (e.g. remote memories, presentations of different stimuli, etc.). A more complete picture of network activity may be provided by unsupervised methods, detecting regularities, for example in the form of spiking patterns recurring more often than predicted by chance. Unsupervised methods proposed so far typically use linear approaches, such as Principal Component Analysis (PCA) (Peyrache et al., 2009; Lopes-dos-Santos, Ribeiro, and Tort, 2013; Stopfer, Jayaraman, and Laurent, 2003), and cannot account for different patterns arising from permutations of spike orderings.

While approaches like frequent itemset mining and related methods (Grun, Diesmann, and Aertsen, 2002; Picado-Muino et al., 2013; Pipa et al., 2008; Torre et al., 2013) can find more patterns than the number of neurons and provide a rigorous statistical framework, they require that exact matches of the same pattern occur, which becomes less and less probable as the number of neurons grows or as the time bins become smaller (problem of combinatorial explosion). To address this problem, Effenberger and Hillar, 2015; Hillar and Effenberger, 2015 proposed another promising unsupervised method based on spin glass Ising models that allows for approximate pattern matching while not being linearly limited in the number of patterns; this method however requires binning, and rather provides a method for classifying the binary network state vector in a small temporal neighbourhood, while not dissociating rate patterns from temporal patterns.

In this paper we introduce a novel spike pattern detection method called SPOTDisClust (Figure 1). We start from the idea that the similarities of two neural patterns can be defined by the trace that they may leave on the synaptic matrix, which in turn is determined by the pairwise cross-correlations between neural activities (Dan and Poo, 2004; Markram et al., 1997). The algorithm is based on constructing an epoch-to-epoch dissimilarity matrix, in which dissimilarity is defined as SPOTDis, making use of techniques from the mathematical theory of optimal transport to define, and efficiently compute, a dissimilarity between two spiking patterns (Monge, 1781; Kantorovich, 1942; Hitchcock, 1941; Rubner, Tomasi, and Guibas, 1998). We then perform unsupervised clustering on the pairwise SPOTDis matrix. SPOTDis measures the similarity of two spike patterns (in two different epochs) by determining the minimum transport cost of transforming their corresponding cross-correlation matrices into each other. This amounts to computing the Earth Mover’s Distance (EMD) for all pairs of neurons and all pairs of epochs (see Methods). Through ground-truth simulations, we show that SPOTDisClust has many desirable properties: It can detect many more patterns than the number of neurons (Figure 2); it can detect complex patterns that would be invisible with latency-based methods (Figures 3–4); it is highly robust to noise, i.e. to the ‘insertion’ of noisy spikes, spike timing jitter, or fluctuations in the firing rate, and its performance grows with the inclusion of more neurons given a constant signal-to-noise ratio (Figure 5); it can detect sequences in the presence of sparse firing (Figure 6); and finally it is insensitive to a global or patterned scaling of the firing rates (Figure 7). We apply SPOTDisClust to V1 Utah array data from the awake macaque monkey, and identify different visual stimulus directions using unsupervised clustering with SPOTDisClust (Figure 8).

**Figure 1.**
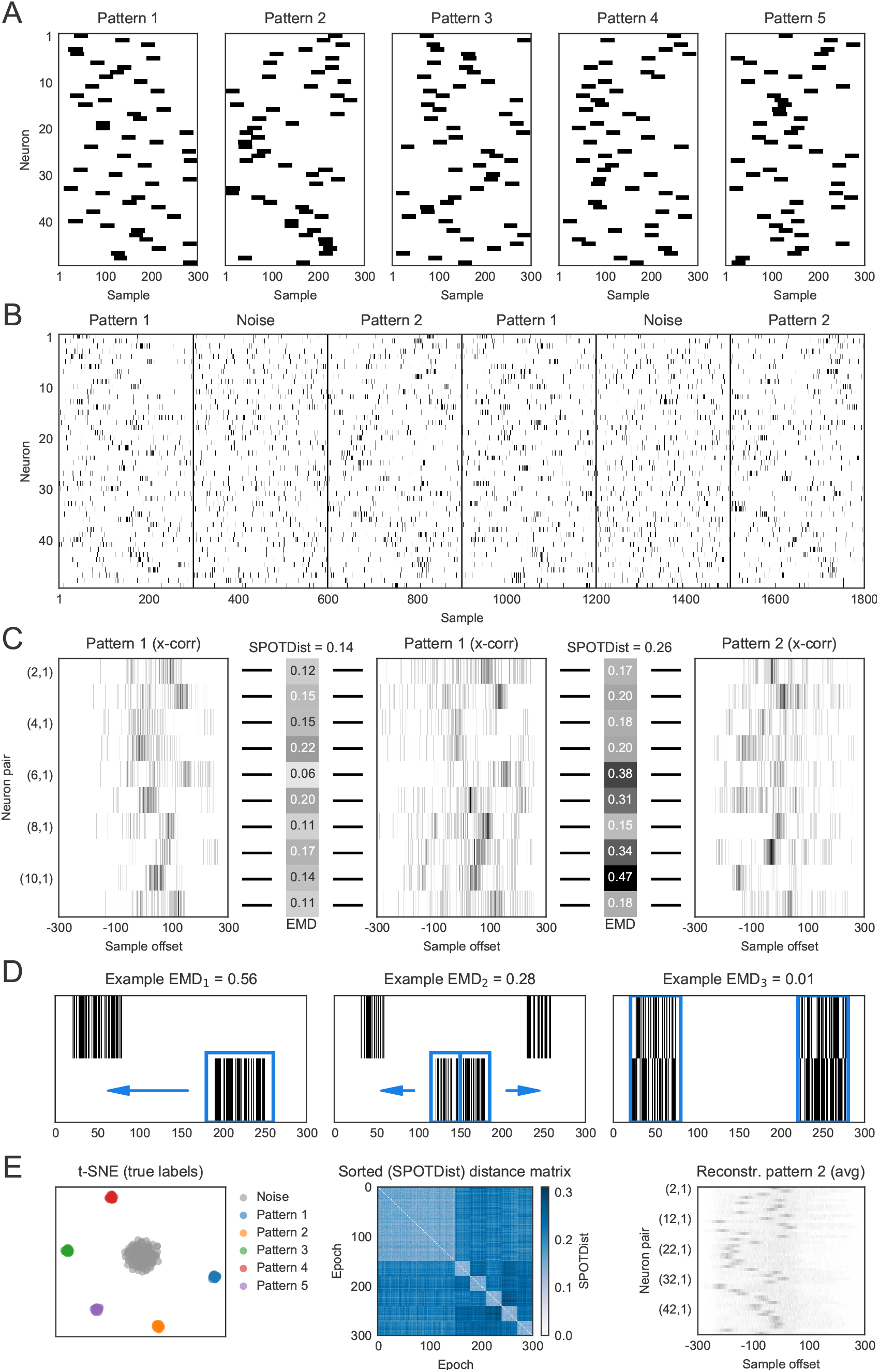
(*preceding page*): Simulated example illustrating the steps in SPOTDisClust. A) Structure of five “ground-truth” patterns, affecting 50 neurons. *λ_in_* = 0.2 spks/sample, *λ_out_* = 0.02 spks/sample. *T_epoch_* = 300 samples, *T_pulse_* = 30 samples. For each pattern and each neuron, a random position was chosen for the activation pulse. B) Neuronal output is generated according to an inhomogeneous Poisson process, with rates dictated by the patterns in (A). A total of 300 epochs were simulated, out of which 150 epochs were noise patterns, and each of the 5 patterns contributed 30 epochs. C) Cross-correlation histograms, normalized to unit mass, are shown for a subset of neurons pairs, for three different epochs. Two epochs constitute realizations of the same pattern, and one epoch belongs to a different pattern. Shown is the EMD for each neuron pair, between the different epochs. These EMDs are then averaged across all neuron pairs to compute the SPOTDis. D) Illustration of the EMD for three different pairs of spike distributions. E) Left: the t-SNE projection with the ground-truth cluster labels shown. Middle: sorted dissimilarity (SPOTDis) matrix, with epochs sorted by the pattern they belong to (first 150 epochs are noise patterns). Right: reconstructed cross-correlations between neuron 1 and neurons 2-49 for pattern 2. Note the similarity between the structure of the reconstructed cross-correlation matrix and the structure of pattern 2.

**Figure 2:**
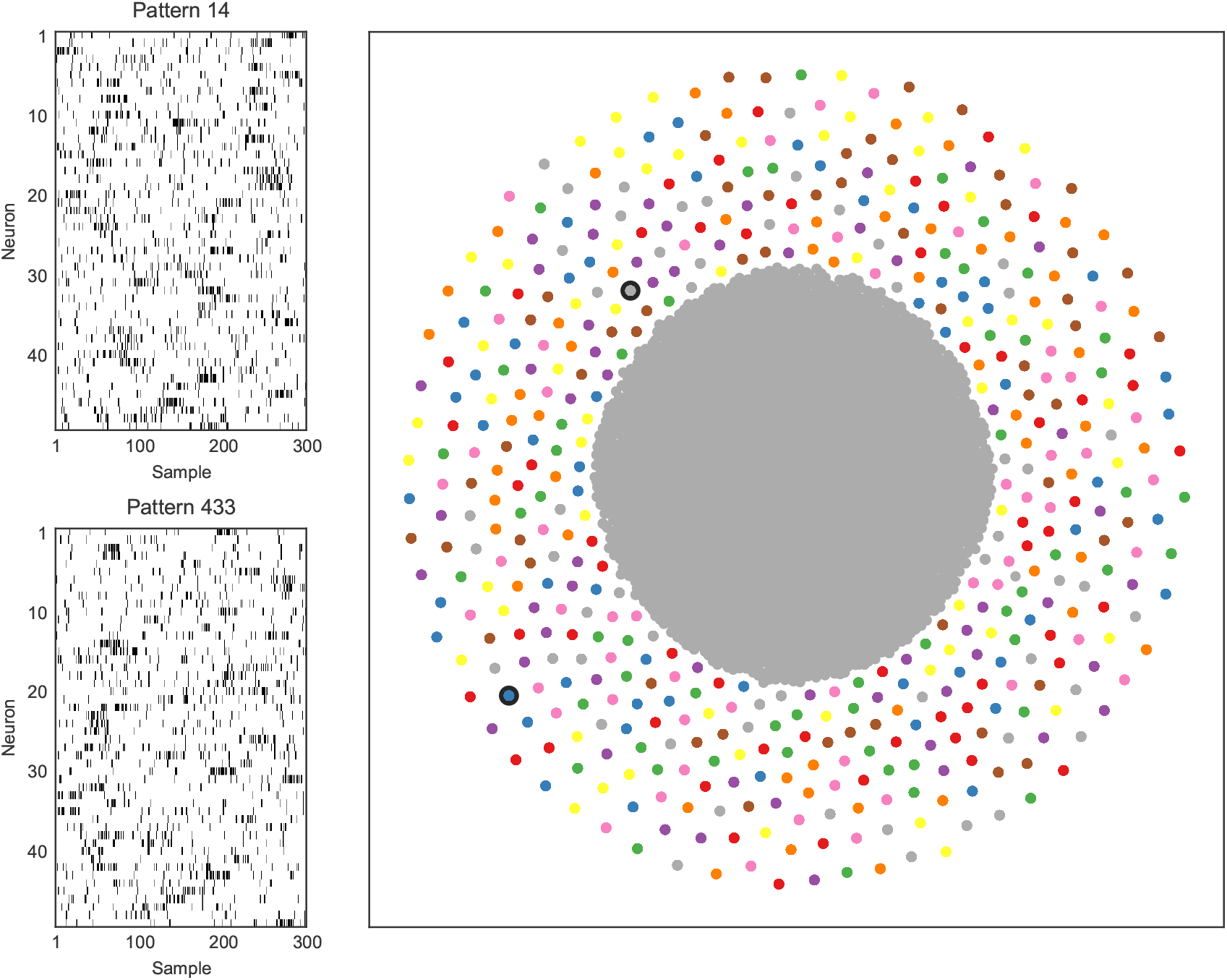
More patterns can be detected than the number of neurons with SPOTDisClust. Left: Two realizations of two different patterns, for 50 neurons. Simulation parameters were *λ_in_* = 0.35 spks/sample, *λ_out_* = 0.05 spks/sample, *T_epoch_* = 300 samples, *T_pulse_* = 30 samples. For each pattern and each neuron, a random position was chosen for the activation pulse. Right: For 500 patterns, 30 realizations per pattern were generated, and 15000 noise epochs were added. t-SNE projection with HDBSCAN labels shows that our clustering method can retrieve all patterns from the data.

**Figure 3.**
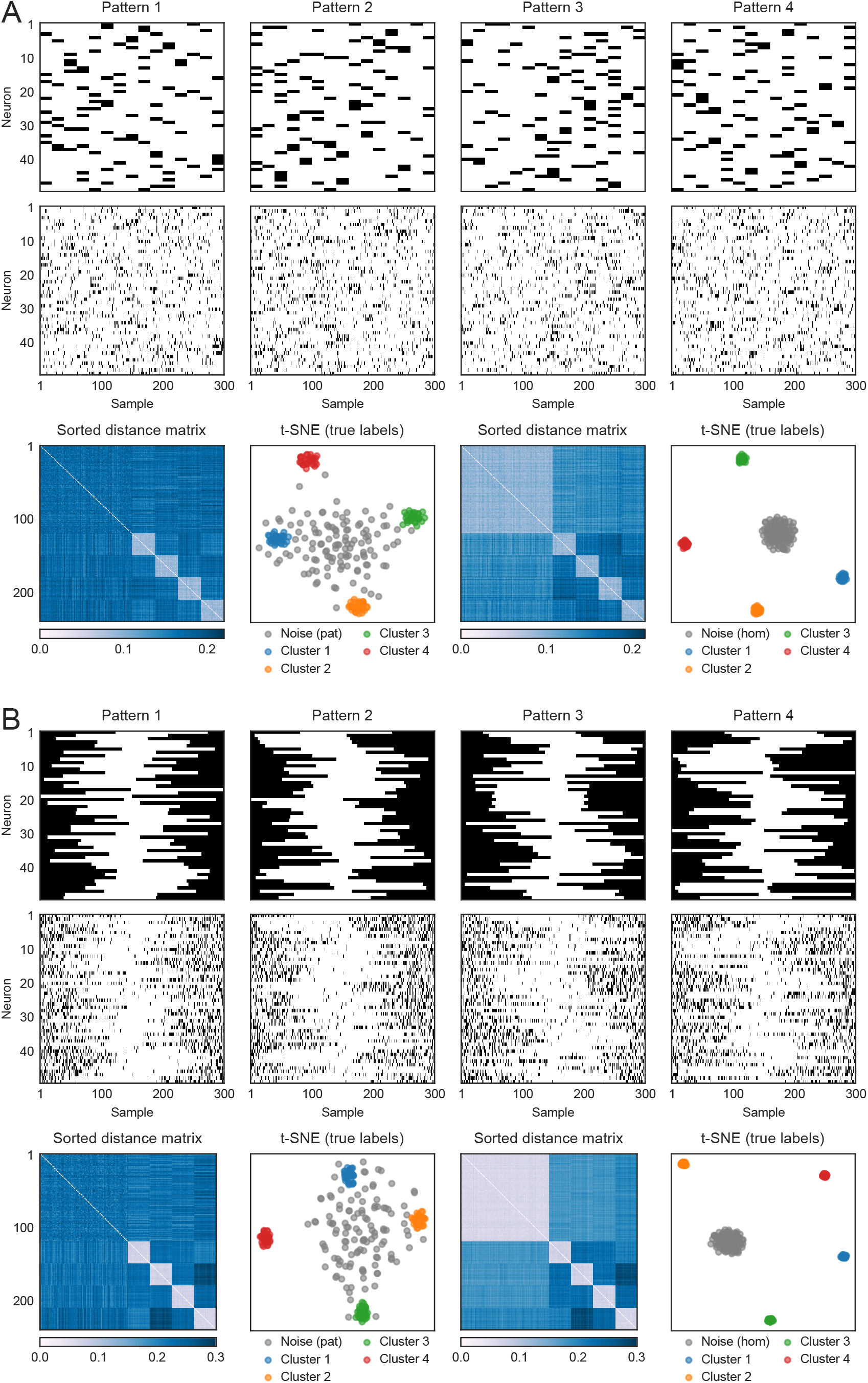
(*preceding page):* Bimodal activation and deactivation patterns can be detected using SPOTDisClust. (A) Multiple bimodal activation patterns and examples of realizations for each pattern (*N* = 50 neurons). Simulation parameters were *λ_in_* = 0.35 spks/sample, *λ_out_* = 0.05 spks/sample, *T_epoch_* = 300 and *T_pulse_* = 20 samples. Bottom figures show sorted dissimilarity matrix and t-SNE for simulation with patterned noise (left) and homogeneous noise (right). (B) Multiple bimodal activation patterns and examples of realizations for each pattern (*N* = 50 neurons). Simulation parameters were *λ_out_* = 0.02 spks/sample (i.e. the deactivation period), λ_in_ = 0.3 spks/sample, *T_epoch_* = 300 and *T_deactivation_* = 150 samples. Bottom figures show sorted dissimilarity matrix and t-SNE for simulation with patterned noise (left) and homogeneous noise (right).

**Figure 4.**
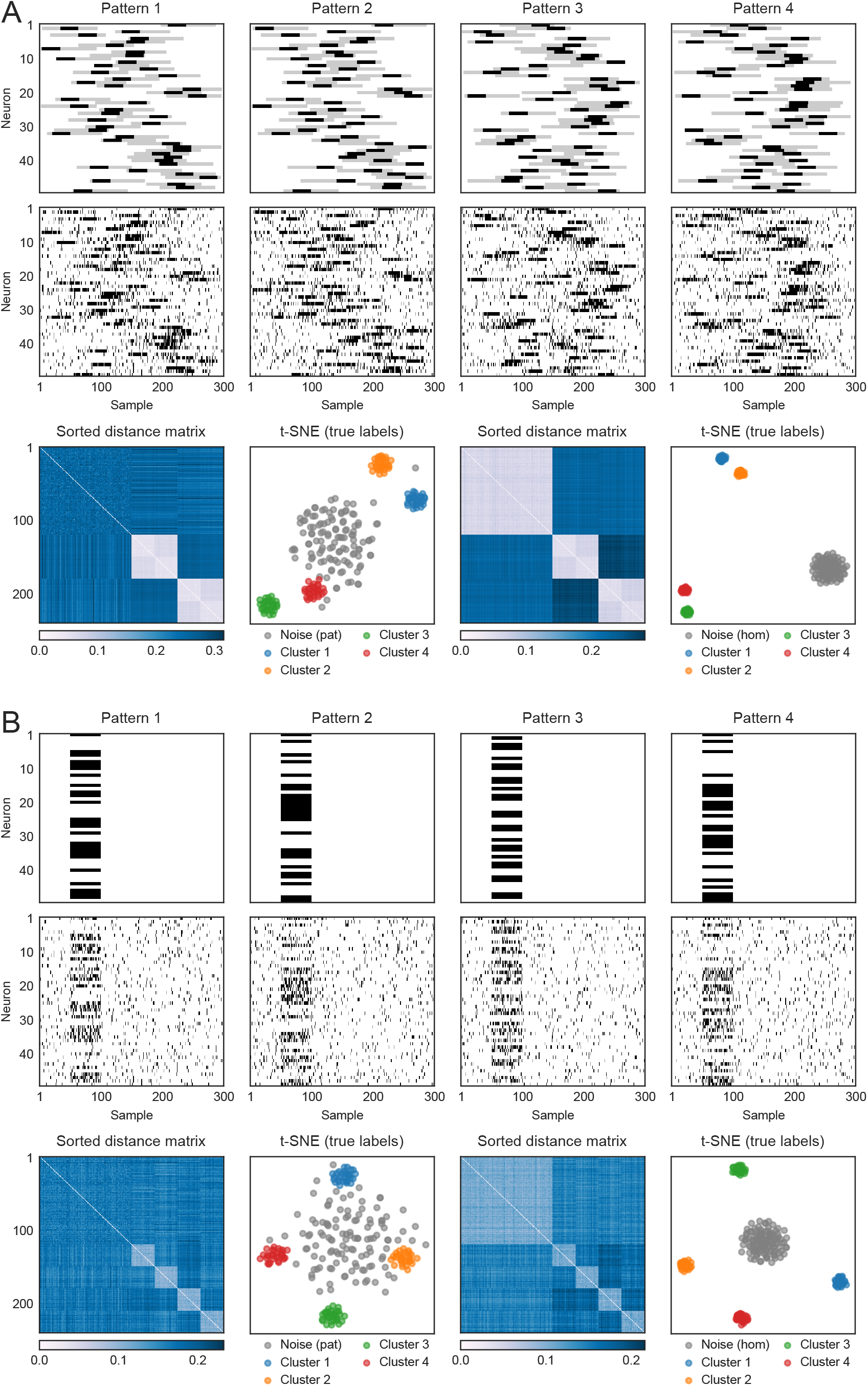
(*preceding page):* Detection of other types of complex patterns using SPOTDisClust. (A) Realizations of four different patterns, in which two patterns (1-2 and 3-4) have the same coarse structure, but a finer is structure embedded inside each coarse pattern. Simulation parameters were *λ_inicoarse_* = 0.2 spks/sample, *λ_in,fine_* = 0.8 spks/sample, *λ_out_* = 0.05 spks/sample *T_epoch_* = 300, *T_pulse,coarse_* = 90 samples, *T_pulse,fine_* = 30 samples. Panels on bottom show sorted dissimilarity matrix and t-SNE for simulations with patterned noise (left) and homogeneous noise (right). (B) Realizations of multiple patterns, in which different random subsets of neurons become simultaneously active, leading to a synchronous firing without temporal order. Simulation parameters were *λ_in_* = 0.4 spks/sample, *λ_out_* = 0.05 spks/sample, *T_epoch_* = 300, *T_pulse_* = 50 samples.

**Figure 5:**
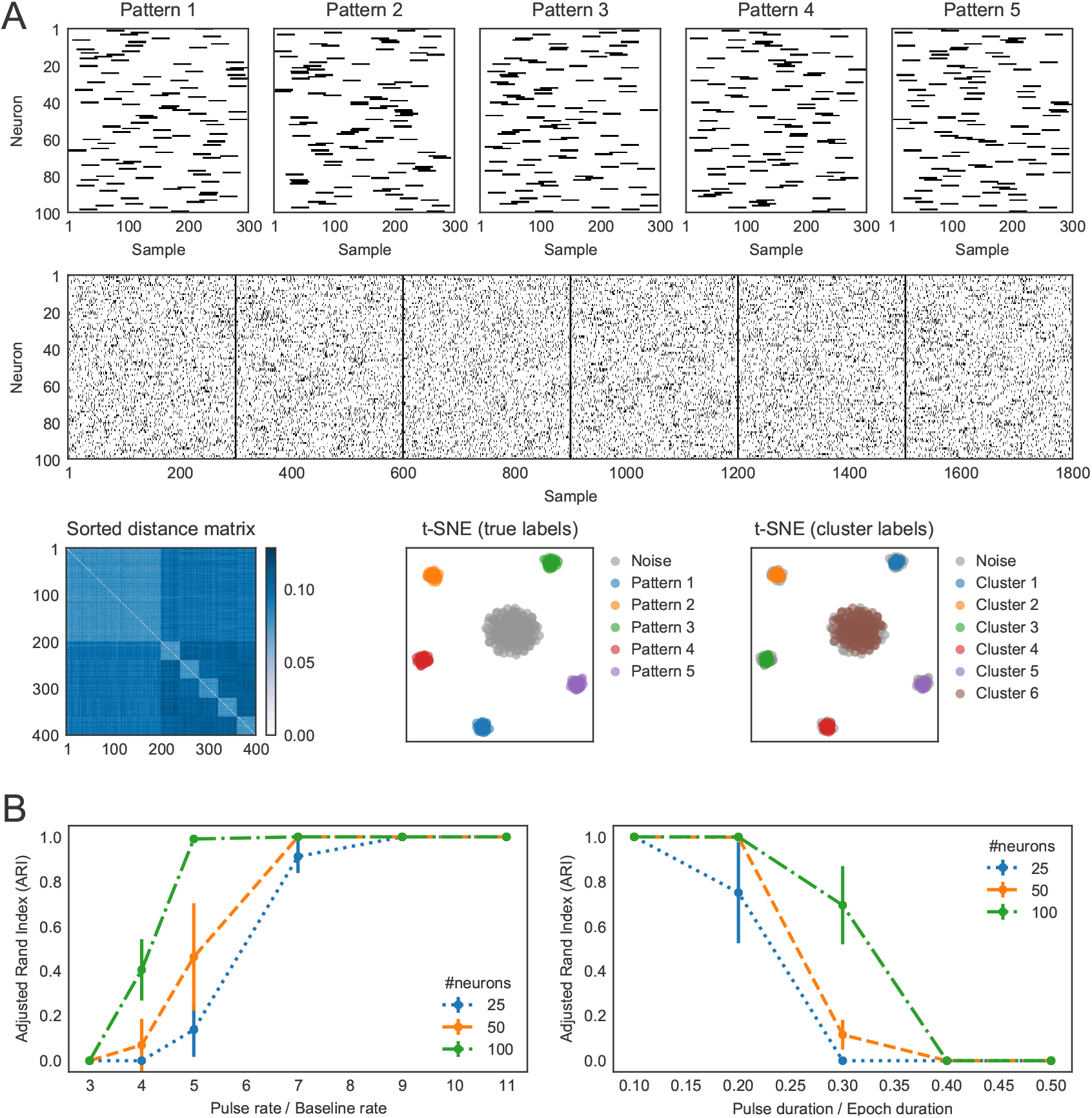
(A) Example of 5 patterns with a low SNR with *N* = 100 neurons and 40 repetitions per pattern, together with 200 noise epochs (homogeneous noise). The order of the spike train realizations is Pattern 1,2, homogeneous noise, 2, 1 and 3. Simulation parameters were *λ_in_* = 0.3 spks/sample, *λ_out_* = 0.1 spks/sample, *T_epoch_* = 300 samples, *T_puise_* = 30 samples. Shown in bottom panels are sorted dissimilarity matrix (left), t-SNE embedding (ground-truth cluster labels), and t-SNE embedding with cluster labels assigned by HDBSCAN. (B) Performance of SPOTDis depends on the SNR. Left: Firing rate inside pulse period is varied, while firing rate outside pulse was varied. We simulated 5 patterns with 30 repetitions each, with *λ_out_* = 0.05 spks/sample, and *λ_in_* attaining values of 0.15, 0.2, 0.25, 0.35, 0.45 or 0.5 spks/sample, *T_pulse_* = 30 and *T_epoch_* = 1000 samples. The number of neurons was 25, 50 or 100. In addition 150 epochs of homogeneous noise were included. We show the mean and the standard deviation across 10 repetitions of the same simulation. Performance relative to ground truth (measured with ARI; see Methods) increases with SNR. Lower SNRs are needed for achieving same performance when the number of neurons is larger. Right: as left, but now varying the pulse duration. Simulation parameters were *λ_out_* = 0.05 spks/sample, and *λ_in_* = 0.5, 0.4, 0.3, 0.2, 0.1 spks/sample, and *T_pulse_* of 100, 200, 300, 400 or 500 samples, with *T_epoch_* = 1000 samples; note that the product of *λ_in_T_pulse_* remained constant.

**Figure 6:**
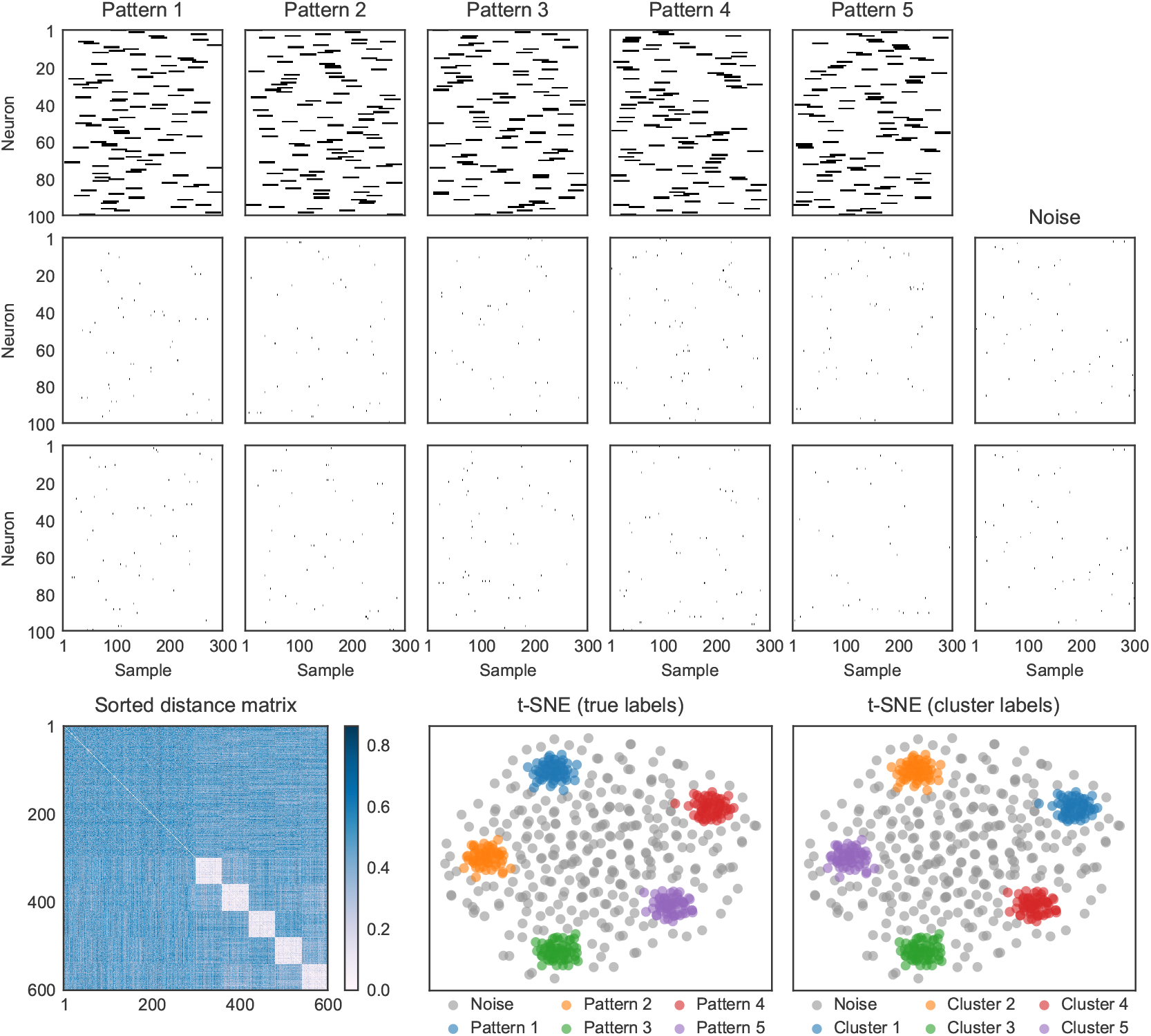
SPOTDis clust can detect temporal patterns expressed in ensembles of sparsely firing neurons. Example of 5 patterns with sparse firing. Simulation parameters were *λ_in_* = 0.015 spks/sample, *λ_out_* = 0.0001 spks/sample, *T_epoch_* = 300 samples, *T_pulse_* = 30 samples. For each pattern, two spike realization are shown. Bottom panels show sorted dissimilarity matrix and t-SNE with ground-truth cluster labels (left) and HDBSCAN cluster labels (right).

**Figure 7.**
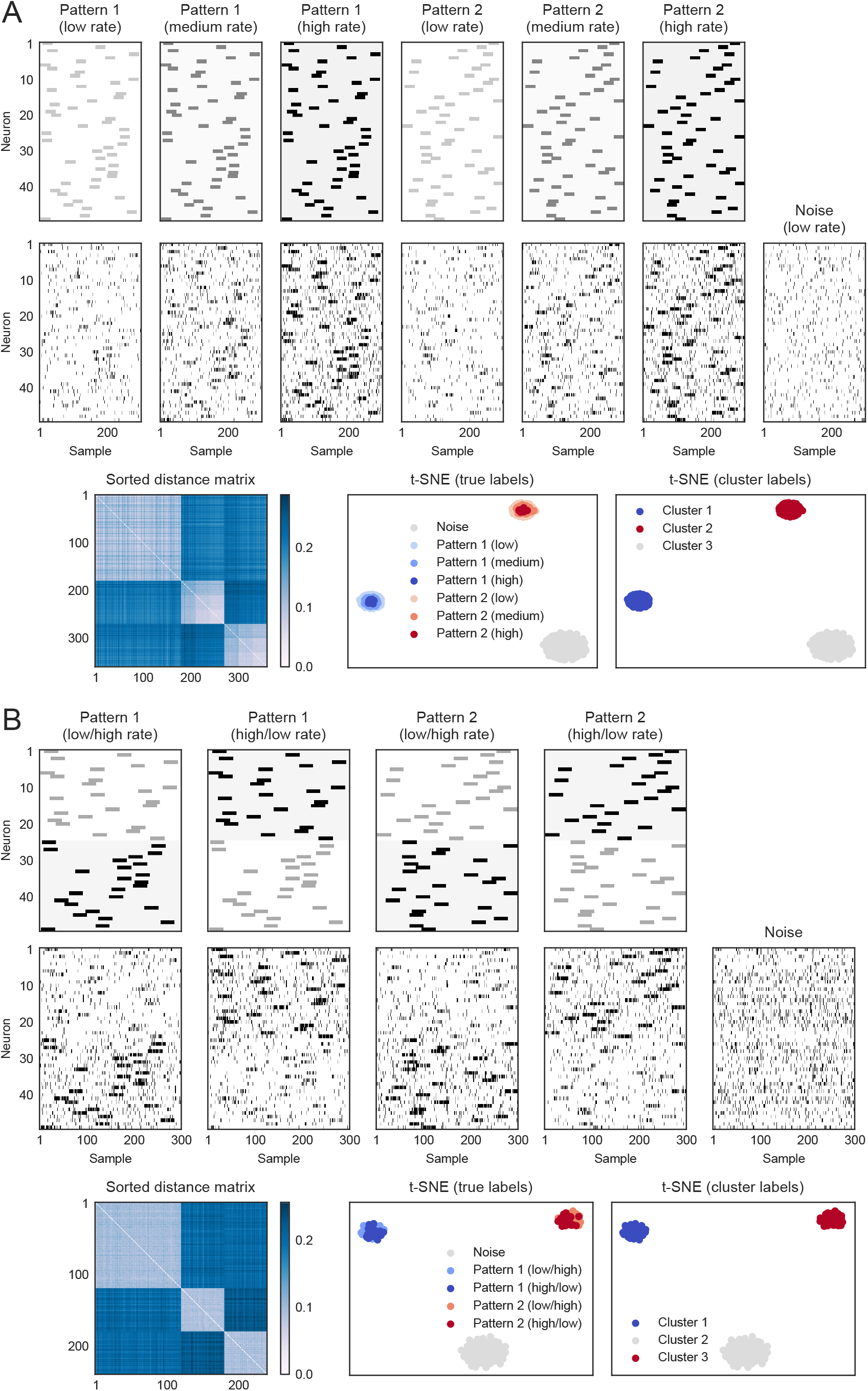
(*preceding page):* Scaling of the rate either globally or in a subset of neurons does not interfere with the detection of temporal patterns. A) Shown are two different temporal patterns. Each temporal pattern can occur in a low (*λ_in_* = 0.2 and *λ_out_* = 0. 02 spks/sample), medium (*λ_in_* = 0.4 and *λ_out_* = 0.04 spks/sample) or high rate (*λ_in_* = 0. 7 and *λ_out_* = 0.07 spks/sample) state, with a constant ratio of *λ_in_/λ_out_.* In addition, the noise pattern can also occur in one of three rate states. The pulse duration was 30 samples. Shown at the bottom the sorted dissimilarity matrix with SPOTDis values, the t-SNE embedding with the ground-truth cluster labels and the t-SNE embedding with the HDBSCAN cluster labels. B) Shown are two temporal patterns. Each temporal pattern could occur in one of two rates states: In the first rate state, the first 25 neurons are firing at low rate (*λ_in_* = 0.3 and *λ_out_* = 0.03 spks/sample), and the other 25 are firing at a high rate (*λ_in_* = 0.7 and *λ_out_* = 0.07 spks/sample). In the second rate state, the rate scaling is reversed. The pulse duration was 30 samples. Shown at the bottom the sorted dissimilarity matrix with SPOTDis values, the t-SNE embedding with the ground-truth cluster labels and the t-SNE embedding with the HDBSCAN cluster labels.

**Figure 8:**
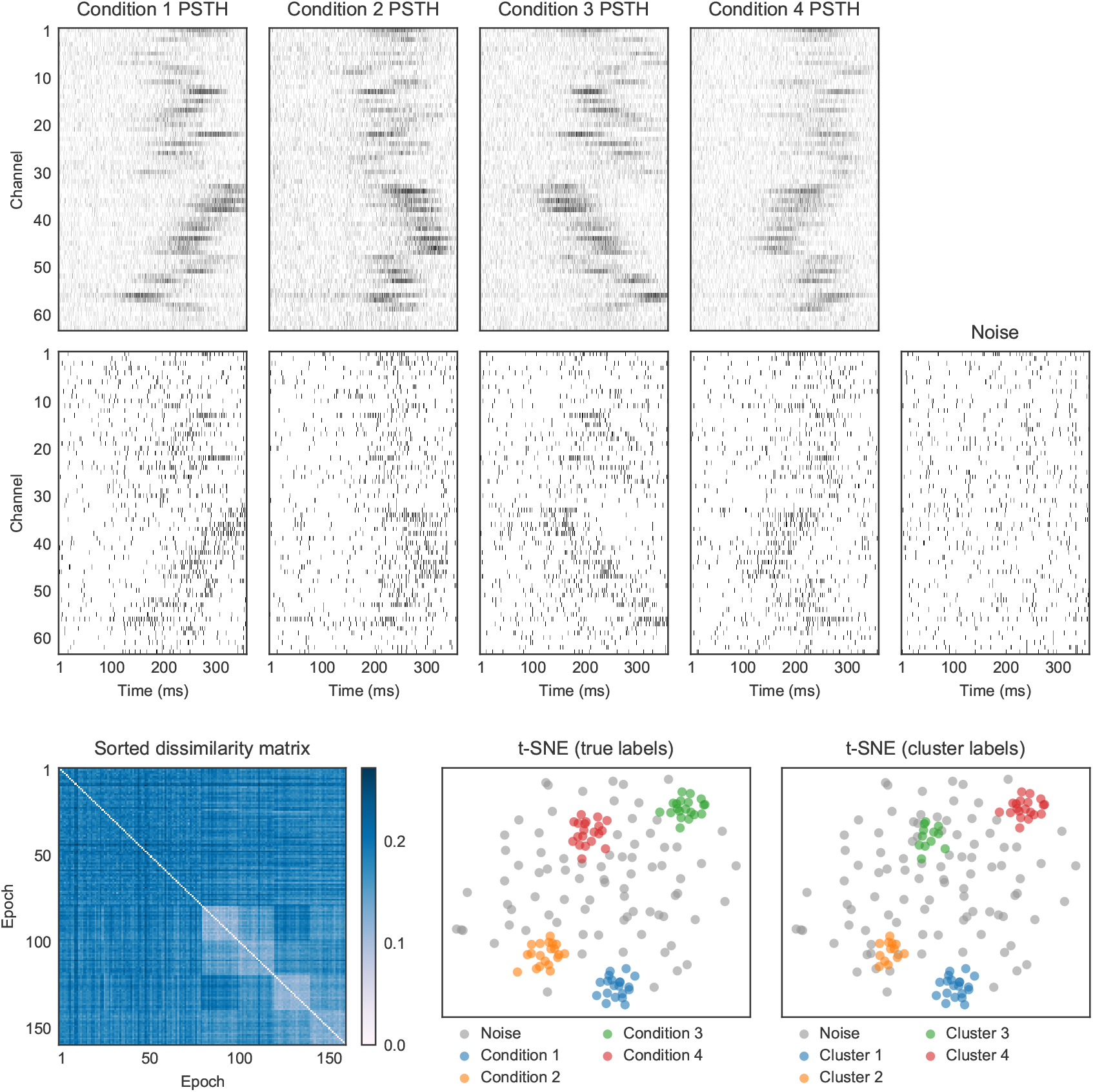
Application to neuronal data. Top: shown are the average peri-stimulus histograms for four moving bar stimuli spanning the four cardinal directions. Middle: multi-unit spike train realizations for each of the four patterns. Bottom: sorted dissimilarity matrix, t-SNE with ground-truth labels, and t-SNE with HDBSCAN cluster labels.

## 2 Results

### 2.1 Outline of the algorithm

Suppose we perform spiking measurements from an ensemble of *N* neurons, and we observe the spiking output of this ensemble in *M* separate epochs of length *T* samples (in units of the time bin length). Suppose that there are *P* distinct activity patterns that tend to reoccur in some of the *M* epochs. Each pattern generates a set of normalized (to unit mass) cross-correlation histograms among all neurons. Instantiations of the same pattern are different because of noise, but will have the same expectation for the cross-correlation histogram. The normalized cross-correlation histogram is defined as

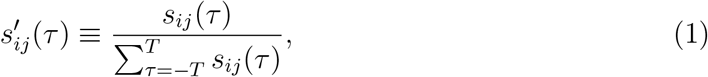

if 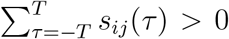, and 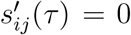 otherwise. Here, 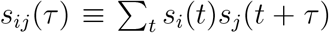 is the cross-correlation function (or cross-covariance), and *s_i_*(*t*) and *β_j_*(*t*) are the spike trains of neurons i and *j*. In other words, the normalized cross-correlation histogram is simply the histogram of coincidence counts at different delays *τ*, normalized to unit mass. We take the *N* × *N* × (2*T* + 1) matrix of 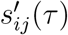 values as a full representation of a pattern, that is, we consider two patterns to have the same temporal structure when all neuron pairs have the same expected value of 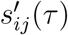 for each *τ*. For simplicity and clarity of presentation, we have written the cross-correlation function as a discrete (histogram) function of time. However, because the SPOTDis, which is introduced below, is a cross-bins dissimilarity measure and requires only to store the precise delays *τ* at which 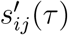 is non-zero, the sampling rate can be made infinitely large (see Methods). In other words, the SPOTDis computation does not entail any loss of timing precision beyond the sampling rate at which the spikes are recorded.

The SPOTDisClust method contains two steps (Figure 1), which are illustrated for five example patterns (Figure 1A-B). The first step is to construct the SPOTDis dissimilarity measure between all pairs of epochs on the matrix of cross-correlations among all neuron pairs. The second step is to perform clustering on the SPOTDis dissimilarity measure using an unsupervised clustering algorithm that operates on a dissimilarity matrix. Many algorithms are available for unsupervised clustering on pairwise dissimilarity matrices. One family of unsupervised clustering methods comprises so called density clustering algorithms, including DBSCAN, HDBSCAN or density peak clustering. Here, we use the HDBSCAN unsupervised clustering method (Ester et al., 1996; Campello, Moulavi, and Sander, 2013; Campello et al., 2015; McInnes, Healy, and Astels, 2017) (see Methods). To examine the separability of the clusters in a low dimensional 2-D embedding, we employ the t-SNE projection method (Maaten and Hinton, 2008; Hinton and Roweis, 2003) (see Methods).

The SPOTDis measure is constructed as follows:

1. We compute, for each of the *M* epochs separately, the cross-correlation function for all pairs of *N* neurons (see Methods), which yields *M* matrices of *N*(*N* – 1)/2 cross-correlations 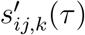 (Figures 1C and S1).
2. For each pair of epochs *k* and *m* and each pair of neurons *i* and *j*, we now want to quantify how similar the temporal correlation of neuron *i* and *j* was between epochs *k* and *m*, i.e. the similarity of 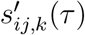 and 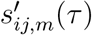. To this end, we compute the Earth Mover’s Distance (EMD) between the normalized cross-correlations of each neuron pair, which yields *M*(*M* – 1)/2 × *N*(*N* – 1)/2 EMD values *D_ij,km_* (see Methods and Figures 1C-D, S1 and S2). We use the L1 norm to measure dissimilarity on the time axis, and we define the cost of transporting the mass in the cross-correlation function between time *τ*_1_ and *τ*_2_ as |*τ*_1_ – *τ*_2_|/(2<u>T</u> + 1), such that the minimum and maximum EMDs are 0 and 1, respectively. The EMD is a metric distance function on probability distributions that determines the minimum transport cost to transform one unit distribution of mass into another unit distribution of mass (Figure 1D). In this case, the mass is the normalized (to a mass of 1) cross-correlation function for a neuron pair. The advantage of using the EMD is multi-fold. First, it is a symmetric and metric measure of similarity between two probability distributions (as opposed to e.g. the Kullback-Leibler divergence). Second, as it is a “cross-bins” distance, it can handle jitter in spike timing. In other words, it quantifies not only whether two distributions are overlapping, like the Kullback-Leibler divergence, but also how far they are shifted away, as minimum transport cost, from each other in a metric space (in this case: time). Third, it does not rely on the computation of a measure of central tendency like the center of mass or peak of the probability distribution, but can also compute transport cost between multimodal probability distributions (Figure 1D). It can therefore capture differences between complex patterns (see Figures 3 and 4). Fourth, because our computation of the EMD uses only the exact pairwise delays among pairs of spikes as its input (the computation thus scales with the number of spikes, not bins; see Methods), our implementation does not require any binning or smoothing of the spike trains beyond the sampling rate at which the spikes are recorded, preventing any additional loss in timing precision; this means that the bins can be made infinitely small (see Methods).
3. After computing the EMDs between each pair of epochs for each neuron pair separately, we compute SPOTDis as the average EMD across neuron pairs,

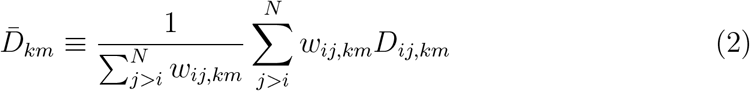

(see Methods) (Figure 1C). Here, the weights are defined as

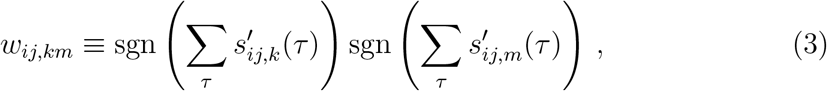

where sgn(*x*) is the sign function, with sgn(*x*) = 0 for *x* = 0 and sgn(*x*) = 1 for *x* > 0. Thus, only for neuron pairs for which both neuron *i* and neuron *j* fired in both epochs *k* and *M* will the weight *w_ij,km_*, = 1. The rationale behind ignoring the other neuron pairs for computing the SPOTDis is that it avoids assigning an arbitrary value to the EMD in the case where we have no information about the temporal relationship between the neurons (i.e. where we do not have any spikes for one neuron in one epoch). We assume for now, that for all 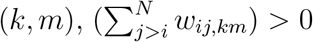, i.e. that for each pair of epochs k and m, there is at least one pair of neurons in which both neurons fired in both epochs k and m. If all the weights equal 1, then we can simplify to

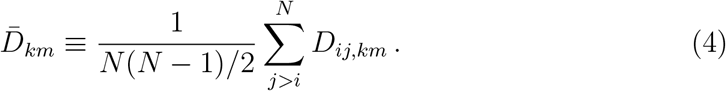 From Eq. (2) we obtain *M*(*M* ‒ 1)/2 SPOTDis values 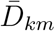 (Figure 1E). These values are then the input to the HDBSCAN clustering algorithm and the t-SNE visualization (Figure 1E) (see Methods).

### 2.2 Ground truth simulations

To test the SPOTDisClust method for cases in which the ground truth is known, we generated *P* input patterns in epochs of length *T* = *T_epoch_* = 300 samples, defined by the instantaneous rate of inhomogeneous Poisson processes, and then generated spiking outputs according to these (Figure 1A-B) (see Methods). Because the SPOTDis is a binless measure, in the sense that it does not require any binning beyond the sampling frequency, the epochs could for example represent spike series of 3s with a sampling rate of 100Hz, or spike series of 300ms with a sampling rate of 1000Hz. Each input pattern was constructed such that it had a baseline firing rate and a pulse activation firing rate, defined as the expected number of spikes per sample. The pulse activation period (with duration *T_pulse_* samples) is the period in the epoch in which the neuron is more active than during the baseline, and the positions of the pulses across neurons define the pattern. For each neuron and pattern, the position of the pulse activation period was randomly chosen. We generated *M*/(2 * *P*) realizations for each of the *P* patterns, and a matching number of M/2 noise epochs (i.e. 50 percent of epochs were noise epochs). We performed simulations for two types of noise epochs (Figure S3). First, noise was generated with random firing according to a homogeneous Poisson process with a constant rate (see Figure 1). We refer to this noise, throughout the text, as “homogeneous noise”. For the second type of noise, each noise epoch comprised a single instantiation of a unique pattern, with randomly chosen positions of the pulse activation periods. We refer to this noise as “patterned noise”. For both types of noise patterns, the expected number of spikes in the noise epoch was the same as during an epoch in which one of the *P* patterns was realized. The second type of noise also had the same inter spike interval statistics for each neuron as the patterns. Importantly, because SPOTDisClust uses only the relative timing of spiking among neurons, rather than the timing of spiking relative to the epoch onset, the exact onset of the epoch does not have to be known with SPOTDis; even though the exact onset of the pattern is known in the simulations presented here, this knowledge was not used in any way for the clustering.

Figure 1 illustrates the different steps of the algorithm for an example of *P* = 5 patterns. For the purpose of illustration, we start with an example comprising five patterns that are relatively easy to spot by eye; later in the manuscript we show examples with a very low signal-to-noise ratio (Figure 5) or sparse firing (Figure 6). We find that in the 2-D t-SNE embedding, the *P* = 5 different patterns form separate clusters (Figure 1E), and that the HDBSCAN algorithm is able to correctly identify the separate clusters (Figure S4). We also show a simulation with many more noise epochs than cluster epochs, which shows that even in such a case t-SNE embedding is able to separate the different clusters (Figure S5). In Figure S3 we compare clustering with homogeneous and patterned noise. The homogeneous noise patterns have a consistently small SPOTDis dissimilarity to each other and are detected as a separate cluster, while the patterned noise epochs have large SPOTDis dissimilarities to each other and do not form a separate cluster, but spread out rather uniformly through the low-dimensional t-SNE embedding (Figure S3).

### 2.3 Detectable patterns outnumber recorded neurons

A key challenge for any pattern detection algorithm is to find a larger number of patterns than the number of measurement variables, assuming that each pattern is observed several times. This is impossible to achieve with traditional linear methods like PCA (Principal Component Analysis), which do not yield more components than the number of neurons (or channels), although decomposition techniques using overcomplete base sets might in principle be able to do so. Other approaches like frequent itemset mining and related methods (Grun, Diesmann, and Aertsen, 2002; Picado-Muino et al., 2013; Pipa et al., 2008) require that exact matches of the same pattern occur.

Because SPOTDisClust clusters patterns based on small SPOTDis dissimilarities, it does not require exact matches of the same pattern to occur, but only that the different instantiations of the same pattern are similar enough to one another, i.e. have SPOTDis values that are small enough, and separate them from other clusters and the noise.

Figure 2 shows an example where the number of patterns exceeds the number of neurons by a factor 10 (500 to 50). In the 2-D t-SNE embedding, the 500 patterns form separate clusters, with the emergence of a noise cluster that has higher variance. Consistent with the low dimensional t-SNE embedding, the HDBSCAN algorithm is able to correctly identify the separate clusters (Figure S4).

When many patterns are detectable, the geometry of the low dimensional t-SNE embedding needs to be interpreted carefully: In this case, all 500 patterns are roughly equidistant to each other, however, there does not exist a 2-D projection in which all 500 clusters are equidistant to each other; this would only occur with a triangle for *P* = 3 patterns. Thus, although the low dimensional t-SNE embedding demonstrates that the clusters are well separated from each other, in the 2-D embedding nearby clusters do not necessarily have smaller SPOTDis dissimilarities than distant clusters when *P* is large.

### 2.4 SPOTDisClust can detect complex patterns

Temporal patterns in neuronal data may consist not only of ordered sequences of activation, but can also have a more complex character. As explained above, a key advantage of the SPOTDis measure is that it computes averages over the EMD, which can distinguish complex patterns beyond patterns that differ only by a measure of central tendency. Indeed, we will demonstrate that SPOTDisClust can detect a wide variety of patterns, for which traditional methods that are based on the relative activation order (sequence) of neurons may not be well equipped.

We first consider a case where the patterns consist of bimodal activations within the epoch (Figure 3A). These type of activation patterns might for example be expected when rodents navigate through a maze, such that enthorinal grid cells or CA1 cells with multiple place fields are activated at multiple locations and time points (O’Keefe and Burgess, 1996; Maurer et al., 2006; Hafting et al., 2005). A special case of a bimodal activation is one where neurons have a high baseline firing rate and are “deactivated” in a certain segment of the epoch (Figure 3B). These kind of deactivations may be important, because e.g. spatial information about an animal’s position in the medial temporal lobe ((Bos et al., 2017)) or visual information in retinal ganglion cells is carried not only by neuronal activations, but also by neuronal deactivations. We find that the different patterns form well separated clusters in the low dimensional t-SNE embedding based on SPOTDis (Figure 3A-B), and that HDBSCAN correctly identifies them (Figure S4).

Next, we consider a case where there are two coarse patterns and two fine patterns embedded within each coarse pattern, resulting in a total number of four patterns. This example might be relevant for sequences that result from cross-frequency theta-gamma coupling, or from the sequential activation of place fields that is accompanied by theta phase sequences on a faster time scale (O’Keefe and Recce, 1993; Dragoi and Buzsaki, 2006). These kinds of patterns would be challenging for methods that rely on binning, because distinguishing the coarse and fine patterns requires coarse and fine binning, respectively. We find that the SPOTDis allows for a correct separation of the data in four clusters corresponding to the four patterns and one noise cluster (Figure 4A), and that HDBSCAN identifies them (Figure S4). As expected, we find that the two patterns that share the same coarse structure (but contain a different fine structure) have smaller dissimilarities to each other in the t-SNE embedding as compared to the patterns that share a different coarse structure.

Finally, we consider a set of patterns consisting of a synchronous (i.e. without delays) firing of a subset of cells, with a cross-correlation function that is symmetric around the delay *τ* = 0 (i.e., correlation without delays). This type of activity may arise for example in a network in which all the coupling coefficients between neurons are symmetric.

Previous methods to identify the co-activation (without consideration of time delays) of different neuronal assemblies relied on PCA (Peyrache et al., 2009), which has the key limitation that it can identify only a small number of patterns (smaller than the number of neurons) Furthermore, while yielding orthonormal, uncorrelated components that explain the most variance in the data, PCA components do not necessarily correspond to neuronal spike patterns that form distinct and separable clusters; e.g. a multivariate Gaussian distribution can yield multiple PCA components that correspond to orthogonal axes explaining most of the data variance.

Figure 4B shows four patterns, in which a subset of cells exhibits a correlated activation without delays. Separate clusters emerge in the t-SNE embedding based on SPOTDis (Figure 4B) and are identified by HDBSCAN (Figure S4). This demonstrates that SPOTDisClust is not only a sequence detection method in the sense that it can detect specific temporal orderings of firing, but can also be used to identify patterns in which specific groups of cells are synchronously co-active without time delays.

### 2.5 Dependence on the signal to noise ratio

A major challenge for the clustering of temporal spiking patterns is the stochasticity of neuronal firing. That is, in neural data, it is extremely unlikely to encounter, in a high dimensional space, a copy of the same pattern exactly twice, or even two instantiations that differ by only a few insertions or deletions of spikes. Furthermore, patterns might be distinct when they span a high-dimensional neural space, even when bivariate correlations among neurons are weak and when the firing of neurons in the activation period is only slightly higher than the baseline firing around it (see further below). The robustness of a sequence detection algorithm to noise is therefore critical.

We can dissociate different aspects of “noise” in temporal spiking patterns. A first source of noise is the stochastic fluctuation in the number of spikes during the pulse activation period and baseline firing period. In the ground-truth simulations presented here, this fluctuation is driven by the generation of spikes according to inhomogeneous Poisson processes. This type of noise causes differences in SPOTDis values between epochs, because of differences in the amount of mass in the pulse activation and baseline period, in combination with the normalization of the cross-correlation histogram. In the extreme case, some neurons may not fire in an given epoch, such that all information about the temporal structure of the pattern is lost. Such a neural “silence” might be prevalent when we search for spiking patterns on a short time scale. We note that fluctuations in the spike count are primarily detrimental to clustering performance because there is baseline firing around the pulse activation period, in other words because “noisy” spikes are inserted at random points in time around the pulse activations. To see this, suppose that the probability that a neuron fires at least one spike during the pulse activation period is close to one for all *M* epochs and all *N* neurons, and that the firing rate during the baseline is zero. In this case, because SPOTDis is based on computing optimal transport between normalized cross-correlation histograms (eq. (5)), the fluctuation in the spike count due to Poisson firing would not drive differences in the SPOTDis.

A second source of noise is the jitter in spike timing. Jitter in spike timing also gives rise to fluctuations in the SPOTDis and in the ground-truth simulations presented here, spike timing jitter is a consequence of the generation of spikes according to Poisson processes. As explained above, because the SPOTDisClust method does not require exact matches of the observed patterns, but is a “cross-bins” dissimilarity measure, it can handle jitter in spike timing well. Again, we can distinguish jitter in spike timing during the baseline firing, and jitter in spike timing during the pulse activation period. The amount of perturbation caused by spike timing jitter during the pulse activation period is a function of the pulse period duration. We will explore the consequences of these different noise sources, namely the amount of baseline firing, the sparsity of firing, and spike timing jitter in Figures 5 and 6.

We define the SNR (Signal-to-Noise-Ratio) as the ratio of the firing rate inside the activation pulse period over the firing rate outside the activation period. This measure of SNR reflects both the amount of firing in the pulse activation period as compared to the baseline period (first source of noise), and the pulse duration as compared to the epoch duration (second source of noise).

We first consider an example of 100 neurons that have a relatively low SNR (Figure 5A). It can be appreciated that different realizations from the same pattern are difficult to identify by eye, and that exact matches for the same pattern, if one would bin the spike trains, would be highly improbable, even for a single pair of two neurons (Figure 5A). Yet, in the 2-D t-SNE embedding based on SPOTDis, the different clusters form well separated “islands” (Figure 5A), and the HDBSCAN clustering algorithm captures them (Figure S4).

To systematically analyze the dependence of clustering performance on the SNR, we varied the SNR by changing the firing rate inside the activation pulse period, while leaving the firing rate outside the activation period as well as the duration of the activation (pulse) period constant. Thus, we varied the first aspect of noise, which is driven by spike count fluctuations. A measure of performance was then constructed by comparing the unsupervised cluster labels rendered by HDBSCAN with the ground-truth cluster labels, using the Adjusted Rand Index (ARI) measure (see Methods). As expected, we find that clustering performance increases with the firing rate SNR (Figure 5B). Importantly, as the number of neurons increases, we find that the same clustering performance can be achieved with a lower SNR (Figure 5B). Thus, SPOTDisClust does not suffer from the problem of combinatorial explosion as the number of neurons that constitute the patterns increases, and, moreover, its performance improves when the number of recorded neurons is higher. The reason underlying this behavior is that each neuron contributes to the separability of the patterns, such that a larger sample of neurons allows each individual neuron to be noisier. This means that, in the brain, very reliable temporal patterns may span high-dimensional neural spaces, even though the bivariate correlations might appear extremely noisy; absence of evidence for temporal coding in low dimensional multi-neuron ensembles should therefore not be taken as evidence for absence of temporal coding in high dimensional multi-neuron ensembles.

We also varied the SNR by changing the pulse duration while leaving the ratio of expected number of spikes in the activation period relative to the baseline constant. The latter was achieved by adjusting the firing rate inside the activation period, such that the product of pulse duration with firing rate in the activation period remained constant, i. e. *T_pulse_λ_pulse_* = *c*. Thus, we varied the second aspect of noise, namely the amount of spike timing jitter in the pulse activation period. We find a similar dependence of clustering performance on the firing rate SNR and the number of neurons (Figure 5B). Hence, patterns that comprise brief activation pulses of very high firing yield, given a constant product *T_pulse_λ_pulse_*, clusters that are better separated than patterns comprising longer activation pulses.

We performed further simulations to study in a more simplified, one-dimensional setting how the SPOTDis depends quantitatively on the insertion of noise spikes outside of the activation pulse periods, which further demonstrates the robustness of the SPOTDis measure to noise (Figure S6).

In addition, we performed simulations to determine the influence of spike sorting errors on the clustering performance. In general, spike sorting errors lead to a reduction in HDBSCAN clustering performance, the extent of which depends on the type of spike sorting error (contamination or collision) (Pillow et al., 2013) and the structure of the spike pattern (Figures S7 and S8). This result is consistent with the dependence of the HBDSCAN clustering performance on signal-to-noise ratio and pulse duration shown in Figure 5, as well with the notion that contamination mixes responses across neurons, such that the number of neurons that carries unique information decreases. We further note that in general, SPOTDisClust provides flexibility when detecting patterns using multiple tetrodes or channels of a laminar silicon probe: A common technique employed when analyzing pairwise correlations (e.g. noise correlations) is to ignore pairs of neurons that were measured from the same tetrode or from nearby channels. When the number of channels is large, this will only ignore a relatively small fraction of neuron pairs. Because the SPOTDis measure is defined over pairs of neurons, rather than the sequential ordering of firing defined over an entire neuronal ensemble, this can be easily implemented by, in Eq. 2, letting the sum run over neuron pairs from separate electrodes.

### 2.6 Temporal pattern recognition in sparsely firing ensembles

As explained above, an extreme case of noise driven by spike count fluctuations is the absence of firing during an epoch. If many neurons remain “silent” in a given epoch, then we can only compute the EMD for a small subset of neuron pairs (eq. (2)). Such a sparse firing scenario might be particularly challenging to latency-based methods, because the latency of cells that do not fire is not defined. We consider a case of sparse firing in Figure 6 where the expected number of spikes per epoch is only 0.48. Despite the firing sparsity, the low-dimensional t-SNE embedding based on SPOTDis shows separable clusters, and HDBSCAN correctly identifies the different clusters (Figure 6). We also performed a simulation in which patterns consist of precise spike sequences, and examined the influence of temporal jitter of the precise spike sequence, as well as the amount of noise spikes surrounding these precise spike sequences. Up to some levels of temporal jitter, and signal-to-noise ratio, HDBSCAN shows a relatively good clustering performance (Figure S9).In general, given sparse firing, a sufficient number of neurons is needed to correctly identify the *P* patterns, but, in addition, the patterns should be distinct on a sufficiently large fraction of neuron pairs. Furthermore, the lower the signal-to-noise ratio, the more neurons are needed to separate the patterns from one another.

### 2.7 Insensitivity to scaling of firing rates

A key aim of the SPOTDisClust methodology is to identify temporal patterns that are based on consistent temporal relationships among neurons. However, in addition to temporal patterns, neuronal populations can also exhibit fluctuations in the firing rate that can be driven by e.g. external input or behavioral state and are superimposed on temporal patterns. A global scaling of the firing rate, or a scaling of the firing rate for a specific assembly, should not constitute a different temporal pattern if the temporal structure of the pattern remains unaltered, i.e. when the normalized cross-correlation function has the same expected value, and should not interfere with the clustering of temporal patterns. This is an important point for practical applications, because it might occur for instance that in specific behavioral states rates are globally scaled (McGinley et al., 2015; Steriade, Timofeev, and Grenier, 2001).

In Figure 7A, we show an example where there are three different global rate scalings, as well as two temporal patterns. The temporal patterns are, for each epoch, randomly accompanied by one of the different global rate scaling factors. The t-SNE embedding shows that the temporal patterns form separate clusters, but that the global rate scalings do not (Figure 7A). Furthermore, HDBSCAN correctly clusters the temporal patterns, but does not find separate clusters for the different rate scalings (Figures 7A and S4). This behavior can be understood from examination of the sorted dissimilarity matrix, in which we can see that epochs with a low rate do not only have a higher SPOTDis to epochs with a high rate, but also to other epochs with a low rate, which prevents them from agglomerating into a separate cluster (Figure 7A); rather the epochs with a low rate tend to cluster at the edges of the cluster, whereas the epochs with a high rate tend to form the core of the cluster (Figure 7A).

Another example of a rate scaling is one that consists of a scaling of the firing rate for one half of the neurons (Figure 7B). Again, the t-SNE embedding and HDBSCAN clustering show that rate scalings do not form separate clusters, and do not interfere with the clustering of the temporal patterns (Figures 7B and S4). We conclude that the unsupervised clustering of different temporal patterns with SPOTDisClust is not compromised by the inclusion of global rate scalings, or the scaling of the rate in a specific subset of neurons.

### 2.8 Application to visual responses of V1 populations

We apply the SPOTDisClust method to data collected from monkey V1. Simultaneous recordings were performed from 64 V1 channels using a chronically implanted Utah array (Blackrock) (see Methods). We presented moving bar stimuli in four cardinal directions while monkeys performed a passive fixation task. Each stimulus bar was presented 20 times. We then pooled all 80 trials together, and added 80 trials containing spontaneous activity. Our aim was then to recover the separate stimulus conditions using unsupervised clustering of multi-unit data with SPOTDisClust. The low dimensional t-SNE embedding shows four dense regions that are well separated from each other and correspond to the four stimuli, and HDBSCAN identifies these four clusters (Figure 8). Furthermore, when we performed clustering on the firing rate vectors, and constructed a t-SNE embedding on distances between population rate vectors, we were not able to extract the different stimulus directions from the cluster labels. This shows that the temporal clustering results are not trivially explained by rate differences across stimulus directions, but also indicates that the temporal pattern of population activity might reveal important stimulus information beyond the neuronal firing rates. Thus, SPOTDisClust can be successfully used on real neuronal data to identify different temporal patterns in high-dimensional multi-neuron ensembles.

### 2.9 Optimization of window length

For the simulations and applications above, we have assumed that the temporal window of interest for the application was known, i.e. that we had some *a priori* knowledge about the length of the spike patterns. For many applications in neuroscience, we want to detect sequences around specific events that are either defined by some external event (e.g. a stimulus onset) or by some “internal” event, e.g. a sharp wave ripple complex, an UP-DOWN state transition, or the cycle of some oscillation, e.g. a hippocampal theta cycle. However, in these cases the precise duration of the spike patterns is not known, and in addition there is most likely some jitter in the onset of the sequence. To handle this onset jitter, SPOTDisClust explicitly defines patterns at the level of cross-correlations rather than on univariate spike trains directly, which endows the measure with time translational invariance. To show the feasability of determining the window length automatically from the data, we have performed simulations in which sequences occurred around some event with additional temporal jitter. These sequences were flanked by homogeneous noise (i.e. before and after the sequences). We then varied the chosen window length around the event, and measured the cluster quality using the unsupervised Silhouette score, which is based on a comparison of distances within each cluster vs. distances between the clusters (see Methods). As expected, the (unsupervised) Silhouette and ARI (ground-truth) cluster quality scores decreased when we used longer and shorter windows than the true sequence length (Figure S11). Importantly, however, the Silhouette and ARI score showed a tight correspondence with one another, showing the feasability of selecting the window length for spike sequence detection in an unsupervised manner. For our application to neuronal data, we optimized the window length used for the clustering using the Silhouette score and show that we can recover the window length that maximizes the cluster quality as compared to ground-truth (ARI) (Figure S12).

## 3 Discussion

We have presented a novel dissimilarity measure for multi-neuron temporal spike patterns, SPOTDis, with unique properties that make it suitable for the unsupervised exploration of the space of admissible firing patterns. SPOTDis is rooted in optimal transport theory, a burgeoning field in mathematics that offers promising solutions for fields as diverse as economics, engineering, physics and chemistry (Monge, 1781; Kantorovich, 1942; Hitchcock, 1941; Rubner, Tomasi, and Guibas, 1998; Villani, 2008). In machine learning, optimal transportation based distances for image classification have been devised, which accommodate the fact that relevant image features may appear at slightly different positions in similar images. While pixel-wise comparisons of two images may fail to recognize similarity under those conditions, optimal transportation based distances operate in a “cross-bins” fashion, so they can treat those shifts in an appropriate way. In neural data analysis we face a similar problem, as spike patterns may present themselves repeatedly with the same overall structure, but not exactly the same timing. The traditional approach to accommodate for such “jitter” is to discretize spike times with a binning procedure, or, in a nearly equivalent way, to use a smoothed version of the spike train time series (van Rossum, 2001). Such approaches require setting an arbitrary scale for the timing precision of neural firing. This is in general difficult, because neural patterns may occur at different temporal scales, and with different jitters. For example, hippocampal place fields fire in sequences at the “behavioral” time scale of hundreds of milliseconds, and because of the phase precession phenomena, they fire so-called “theta” sequences at a much faster (tens of milliseconds) time scales (Dragoi and Buzsaki, 2006; O’Keefe and Recce, 1993; Pastalkova et al., 2008). Repeated sequences at any time scale will be detected by SPOTDis, in particular in combination with a density based algorithm such as HDBSCAN, which can detect state space regions of higher density surrounded by lower region areas, regardless of the absolute density. Using ground-truth simulations, we have shown that SPOTDis can deal with cases in which both coarse and fine patterns co-exist (Figure 4A). Optimal transport theory provides both theoretical grounding, as well as a host of solutions for the efficient calculation of distances. Here, we propose a novel implementation, inspired to work in optimal transport, and tailored to the case of calculating the dissimilarity between point process realization, in our case spike trains.

Distance measures based on “morphing” one spike train into another by moving spikes have been previously proposed. The Victor-Purpura distance, which is an adaptation of the Levenshtein distance to point processes, is a paradigmatic example (Victor and Purpura, 1996). Our approach differs in two fundamental ways.

First, the Victor-Purpura distance allows for the insertion and deletion of spikes, to enable computation of distances between spike trains with different numbers of spikes, adding in each case a penalty term (the penalty terms are arbitrary parameter to be optimized). While this may be a principled way to deal with this issue, it introduces additional complexity in the computation of the distance as many different combinations of spike shifting and insertion/deletion must be considered in order to find the optimal solution. This may render optimization difficult and the computation prohibitive as one attempts to compare a large number of multi-neuron patterns. We take the more simple-minded approach of rescaling the time series to be compared, in order to equalize mass. While this may be an oversimplification in some cases, this enables us to implement the computation in a very efficient way. Yet, we preserve many desirable features of spike train metrics such as the Victor-Purpura distance. For example, SPOTDis is not based on measures of central tendency, but can also compute dissimilarities between multimodal probability distributions (Figure 3A-B). Furthermore, SPOTDis is particularly noise robust, because it can handle jitter in spike timing, as it does not require exact overlap in discretized time bins, but is based on distance computations in a metric space.

A second important difference with spike train metric methods such as Victor-Purpura distance is that we calculate the pairwise epoch-to-epoch dissimilarity not directly on spike trains but on cross-correlograms between pairs of cell spike trains. This has the considerable advantage of enabling detection of similarity between spiking patterns that are misaligned, and eliminates the need for precise time reference points (e.g. the time of stimulus delivery), providing a way to freely search for repeated patterns in spontaneous or evoked activity. Comparing cross-correlation patterns between epochs has been used in seminal work on memory replay, where cross-correlation “bias” was compared across entire sleep or behavioral epochs, to assess the presence of significant replay (Skaggs and McNaughton, 1996; Euston, Tatsuno, and McNaughton, 2007). Here, we provide a method for comparison at a greater granularity, enabling efficient identification of the repeated patterns within time windows of hundreds of milliseconds. A distance based on cross-correlation also has a attractive physiological interpretation: From the perspective of synaptic plasticity, it can be interpreted as the extent to which two patterns have a similar effect on the synaptic plasticity in the network through the STDP rule, which holds that changes in synaptic plasticity depend on the timing jitter between pre- and post-synaptic spikes (Dan and Poo, 2004; Markram et al., 1997). Our dissimilarity measure acts on multi-neuron patterns, and can make use of any additional information available when the monitored neural population increases in size. Because SPOTDis between epoch k and epoch *M* ignores neuron pairs in which one neuron did not fire in both epochs, it also handles cases in which there is sparse firing and many neurons do not fire at all (Figure 6). Moreover, SPOTDis is based on computing a distance function on the normalized cross-correlation functions. Because of this normalization to unit mass, it copes with global fluctuations in the firing rate, and specific increases in the firing rate for subsets of neurons (Figure 5).

We combined SPOTDis with a density-based clustering algorithm, HDBSCAN, which forms a good match for several reasons: First, it can deal with non-metric dissimilarities. While SPOTDis on a single cell pair cross-correlation is metric (and the sum of metrics is a metric), absence of firing in some neurons and in some cell pairs may cause violation of metricity, which is handled gracefully by HDBSCAN. Second, it can identify clusters at different characteristic densities in different regions of the state space, adapting to patterns that may arise at different time scales and different precision due to disparate underlying mechanisms. Yet, other clustering strategies than HDBSCAN may work successfully as well. We show that in many cases, a non-linear embedding technique such as t-SNE acting on SPOTDis yields a quite intuitive representation of the underlying structure of the data.

It is important to emphasize that our method is an explicit clustering method, that can find unique patterns of network activity that are well separated from one another. Several methods using decomposition techniques like PCA or matrix factorization have been utilized with the goal of extracting patterns or sequences from neuronal ensemble data (Peyrache et al., 2009; Mackevicius et al., 2018; Lopes-dos-Santos, Ribeiro, and Tort, 2013; Stopfer, Jayaraman, and Laurent, 2003). We highlight several differences between SPOTDisClust and decomposition techniques: 1) In principle, decomposition techniques like PCA achieve a different goal, namely to identify components that explain a large fraction of variance in the data. These components might in some cases correspond to separate patterns, but do not necessarily so: For instance, a multivariate Gaussian model distribution may yield few components explaining most variance, but these components do not necessarily correspond to clusters. Further, decomposition techniques like PCA decompose into an orthogonal base set, while different spike patterns may in fact be strongly correlated to one another. Thus, decomposition and clustering techniques can provide complementary information. 2) SPOTDisClust can also handle cases where there is a very large number of noise patterns and only few realizations of spike patterns yielding small clusters. These small clusters might drive only a small degree of variance, and may be invisible to decomposition techniques. 3) Many decomposition techniques are designed to find a few components corresponding to dominant axes of variance in the data, yielding fewer “spike patterns” than the number of neurons, although decomposition techniques using overcomplete base sets may in principle find more patterns than the number of neurons. SPOTDisClust on the other hand can find many more patterns than the number of clusters. 4) Finally, SPOTDisClust is in principle a “binless” method, while decomposition techniques will typically require some form of binning resulting in information loss.

We provide an initial application of the SPOTDis measure to real neuronal data, by analyzing multi-electrode recordings in visual cortex. In this analysis, we fed the algorithm the neural data without any knowledge of the task structure, or of the times of stimulus delivery. Strikingly, the identified clusters faithfully reflected the structure of the PSTH calculated with traditional methods, with availability of the stimulus delivery times and labels. Thus, we can recover stimulus information even after normalizing away firing rate information, which is conventionally used to decode different stimuli, demonstrating that the temporal structure of population activity encodes different moving stimulus directions. We also developed an analogous clustering method to SPOTDisClust by constructing a dissimilarity matrix based on L1 distances among population firing rate vectors. Using this clustering technique, the different stimulus directions could not be separated from one another using t-SNE embedding or HDBSCAN.

While we argued that our approach using bivariate cross-correlations yields many advantages, it also has a limitation in the sense that it does not capture higher-order correlations among neurons. Future extensions of this technique may explicitly construct a dissimilarity measure based on high-order correlations among neurons. Indeed, incorporating this may be an interesting avenue for future work. Nonetheless, it should be noted that higher order correlations in a population may be captured by models fitting the first and second moments alone (Tavoni et al., 2017; Meshulam et al., 2017)).

In the present work, we only considered spike trains as if they were recorded using electrophysiological methods. However, this method may also be applied to two-photon calcium imaging data using sensors like GCaMP6f. Analysis of this type of data always involves some additional preprocessing steps like denoising, deconvolution, region of interest identification, and normalization. An excellent strategy would be to apply the method on calcium imaging data after deconvolution and source extraction, which yields sparse time series with “spikes”, although not measured with the same temporal resolution as in case of electrophysiological recordings (Pachitariu et al., 2016; Pnevmatikakis et al., 2016). After deconvolution, the application of SPOTDisClust is straightforward. If one operates on the raw fluorescence data, using a defined set of ROIs corresponding to e.g. somata or spines, then one would have to perform some normalization to get rid of background fluorescence. One possible normalization could be *ΔF* = *F* – *F_background_*, where *F_background_* is the background fluorescence. After this, one would normalize ΔF to unit mass, and then directly compute the EMD on this unit mass.

In conclusion, we have proposed a new tool for the efficient unsupervised analysis of multi-neuron data, which opens up more flexible ways to analyze spontaneous and evoked activity than it has been so far possible.

## 4 Methods

### 4.1 Construction of SPOTDis

The SPOTDisClust method contains two steps. The first step is to construct the pairwise epoch-to-epoch SPOTDis measure on the matrix of cross-correlations among all neuron pairs.

1. We compute, for each of the *M* epochs separately, the normalized cross-correlation function for all pairs of *N* neurons, which yields *M* matrices of *N*(*N* –1)/2 normalized cross-correlations 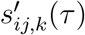. This function is defined

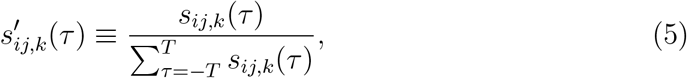

where 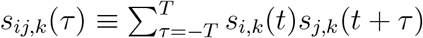 is the cross-correlation function (or cross-covariance), and *s_i,k_* (*t*) and *s_j,k_* (*t*) are the spike trains of neurons *i* and *j* in the kth trial, with *k* = 1,…, *M*, and *i,j* = 1,…, *N*. In other words, the normalized cross-correlation histogram is simply the histogram of expected coincidence counts at different delays *τ*, normalized to unit mass. We define 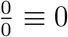 in eq. (5). Figures S1 and S2 show an example of cross-correlations before and after normalization.
2. We then compute the Earth Mover’s Distance (EMD) between 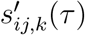 and 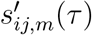 for each (*k, m*). While the EMD is usually defined on each entry of the histogram (i.e. for each *τ*), we can ignore the values of *τ* for which 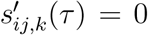 here since there is no mass to move to or from. We therefore first find all the *τ* for which *s_ij,k_*(*τ*) > 0 and *s_ij,m_*(*τ*) > 0, defining respectively the (in ascending order) sorted vectors 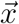 and 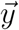 (we omit *i* and *j* subscripts here). These have 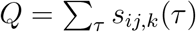 and 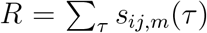 elements respectively, and associated mass 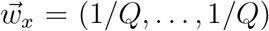 and 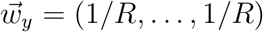. If there exist *τ* for which 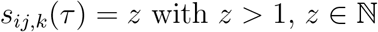, then there will be *z* elements of *τ* in 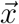 and associated mass in 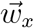, the same for *m* and 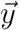. Following these definitions, we have *s_ij,k_*(*x_q_*) > 0 for all *q* and *s_ij,m_*(*y_r_*) > 0 for all *r*, i.e. the vectors 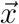 and 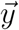 contain all the pairwise delays between the spike trains of a pair of neurons in epoch *k* and *m*, respectively. Furthermore, *x_q_* ≥ *x_p_* for all *q* > *p*, and and *y_r_ ≥ y_z_* for all *r* > *z*, i.e. the vectors contain pairwise delays in ascending order. As an example, if neuron *i* fired spikes at samples (10,11, 20, 23) in epoch *k* and neuron *j* fired spikes at (14,15, 20) then 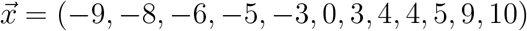 and 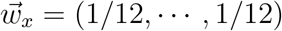. Because the EMD is computed on the precise delay times, we can let Δ*t* → 0 and Δτ → 0, i.e. the sampling rate can be made infinitely large. In practice, we therefore directly find the vectors 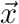 and 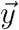 and do not compute 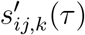 for all 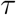. The EMD also requires the definition of a moving cost function, which in this case is defined over time points. Let *c* be the moving cost function which we define as the L1 norm, *c*(*τ*_1_, *τ*_2_) |*τ*_1_ – *t*_2_|/(2*T* + 1). Note that the normalization of 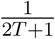 ensures that 0 ≤ *D* ≤ 1 (the EMD) in eq. (6). Solving the optimal transport problem then amounts to finding a matrix of flows **F** ≡ [*f_q,r_*], with *f_q,r_* the flow (i.e. the amount of mass moved) from *w_q,x_* to *w_r,y_*, such that the overall cost is minimized, i.e.

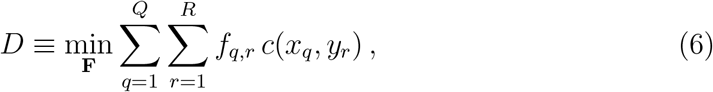

subject to the constraints that

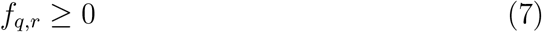

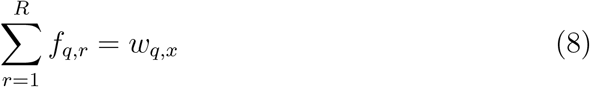

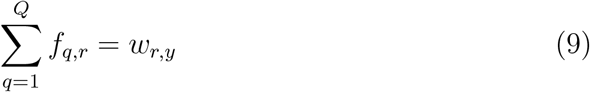

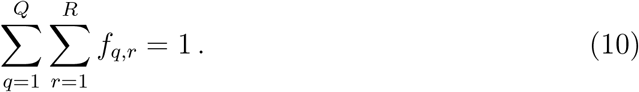 The last equation (10) ensures that all mass needs to be moved (the total mass equals 1), equations (8) and (9) ensure that all the mass from 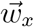 and 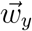 gets moved, and the first equation ensures that the flow is greater or equal than zero. Here, *D* is the EMD. This follows the standard definition of the EMD with normalized mass. Note that the EMD with the L1 norm is equivalent to the 1st order Wasserstein distance (which is a special case of the Kantorovich formulation of the optimal transport). However, for simplicity, we use the notation of the EMD here, which uses discrete variables. We solve the transport problem algorithmically as follows (Figure S2). The algorithm can be intuitively interpreted (as reflected in the comments in pseudo-code) as transporting mass from 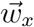 to 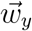, in case *Q* > *R*, with cost *c*(*x_q_,y_r_*). % START ALGORITHM SET emd = 0, *q* = 0, *r* = 0 WHILE *q* < *Q* DO % Move all the remaining mass in *w_q,x_* to *w_r,y_*, but do not move more than *w_r,y_*. SET flow = min(*w_q,x_*, *w_r,y_*) % The *cost* equals the amount of mast moved, *flow*, times the cost of moving % from the time point *x_q_* to the time point *y_r_*. SET cost = flow × *c*(*x_q_,y_r_*) % Add this cost to the total cost. SET emd = emd + cost % Compute the remaining mass in *w_q,x_* to be moved, which equals the % previous mass *w_q,x_* minus the mass we just moved, *flow.* SET *w_q,x_* = *w_q,x_* - *flow* % Compute the remaining mass in *w_r,y_*, to which we still need to transport from 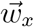 in the next iterations. SET *w_r,y_* = *w_r,y_ – flow* % If there is no mass left then increment the index, note that the vectors 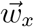 and 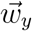 are sorted. IF *w_r,y_* = 0 THEN SET r = r + 1 ENDIF IF *w_q,x_* = 0 THEN SET q = q + 1 ENDIF ENDWHILE % END ALGORITHM
3. After computing the EMDs between each pair of epochs for each neuron pair separately, with computational complexity of order 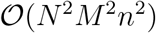, where *n* is the average number of spikes, we compute SPOTDis as the average EMD, i.e.

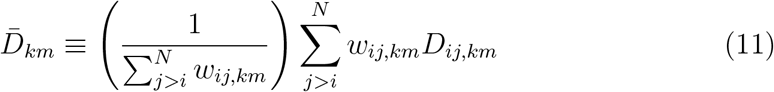

with

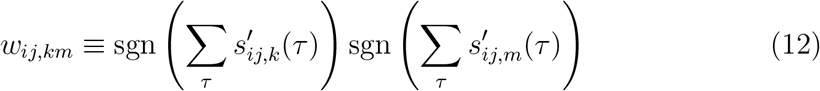

where sgn is the sign function.

Computing the SPOTDis with the sparse simulation (Figure 6) takes 60 seconds and the computation for the example case (Figure 1) takes approximately 5 minutes, utilizing an Intel Xenon E5-2650 v2 2.60GHz system with 16 cores.

### 4.2 HDBSCAN

HDBSCAN is an automated density clustering algorithm that clusters on the basis of pairwise dissimilarity matrices. An extensive overview of HDBSCAN can be found in (Campello et al., 2015; McInnes, Healy, and Astels, 2017) and we provide only a brief overview of HDBSCAN here. HDBSCAN comprises the following steps:

1. After pairwise distances have been computed between all data points, HDBSCAN defines a “mutual reachability distance” between each pair of data points (in our case epochs). The mutual reachability distance is an adjustment of the distance measure that effectively acts as a smoother. For each epoch *k*, the core distance 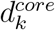 is defined as the SPOTDis dissimilarity, 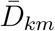 (eq. (2)), to its *n_pts_*th nearest neighbour *m*. The mutual reachability distance is then defined as 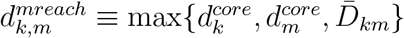. The mutual reachability distance does not alter the distance between two points that are in dense regions, but it changes the distance for points that are in low-density regions and have a relatively large 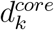 distance. The purpose of transforming the distance matrix in this way is to make the clustering algorithm more robust to noise.
2. HDBSCAN then defines a minimum spanning tree, in which there is a path between all points (vertices), without any loops (i.e. an acyclic graph), such that the total weight of the edge connections is minimized. Here the edges are the mutual reachability distances.
3. HDBSCAN then constructs an hierarchical cluster dendrogram from the minimum spanning tree as follows: Initially all points are assigned to the “root”, a single cluster containing all points. HDBSCAN then sets a threshold *ϵ* and cuts edges from the minimum spanning tree whose weight is higher than *ϵ*. HDBSCAN keeps decreasing the value of *ϵ* such that more connections are removed and new clusters can appear, forming a cluster dendrogram. Sets of points that have fewer than *n_clSize_* members, the minimum cluster size, at a value of *ϵ* are deemed noise points at that value of *ϵ*. Here we take the simplification *n_pts_* = *n_clSize_* (Campello et al., 2015), such that *n_pts_* is the only hyperparameter, which we set to 10 here, unless specified otherwise. We use the implementation of HDBSCAN developed by McInnes, Healy, and Astels (2017), and all analyses and simulations were performed in Python. HDBSCAN uses either the “leaf selection” method for selecting clusters, or the “excess of mass” method. In the leaf selection method, the end branches of the hierarchical cluster dendrogram are taken as the selected clusters. In the “excess of mass” algorithm, HDBSCAN chooses the set of clusters that is most stable under a change of *ϵ*. This is done as follows: at some value of *ϵ* a cluster is born, *ϵ_max_*, and at some point, the cluster dies, *ϵ_min_*. For each member of the cluster we can define the value of *ϵ_k_* where each kth member fell out, and take the stability as the sum 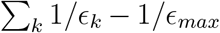 over all members. HDBSCAN then selects in the dendrogram the optimal levels at which to cut the tree in order to maximize the stability of the selected clusters, forming a set of clusters. An advantage of this selection procedure is that it allows for clusters of varying density.

### 4.3 t-SNE

T-SNE (t-distributed stochastic neighbor embedding) is a dimensionality reduction technique for high-dimensional datasets (Maaten and Hinton, 2008). While it typically is computed starting from a high-dimensional dataset that is then converted into a matrix of pairwise Euclidean distances, here we compute it directly on the pairwise dissimilarity matrix. We first outline the algorithm of SNE (Hinton and Roweis, 2003), and after that the adjustments made in t-SNE.

1. For each two data points (*k* and *m*), which represent epochs in our case, a measure of similarity is computed. This measure of similarity is taken as the conditional probability of observing *m* given a Gaussian distribution centered on *k*,

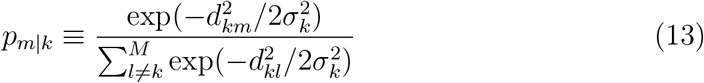 Here, the standard choice for *d_km_* is the L1 norm for a high dimensional dataset *x*_1_,…, *x_M_*, defined as *d_km_* = ||*x_k_* – *x_m_*||, but we take it here as the SPOTDis, 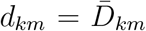. The normalization in eq. (13) simply assures that the probabilities Σ_*m*_*P_m|k_* sum to one. The variance of the Gaussian, 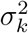, is determined for each data point individually, by finding *σ_k_* to satisfy the equation

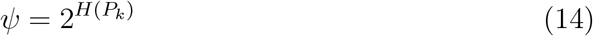 Here, *H* is the entropy function using logarithm base 2, *P_k_* is the probability distribution of all the data points given *k, P_k_* = (*p_1|k_*,…,*p_M|k_*), and *ψ* is the perplexity which is set by the user, which can be interpreted as a smooth measure of the effective number of neighbors each datapoint has. We set the perplexity *ψ* to 30, which is in the typical range used in the literature. T-SNE is generally quite insensitive to choices of the perplexity, which is usually taken in the range 5-50.
2. SNE then attempts to find a low dimensional set of data points, {y_1_,…, y_*M*_}, that have a similar distribution of conditional probabilities (similarities) as the distances derived from the high-dimensional counterparts. In this case the variance of the Gaussian is constant for all data points, and

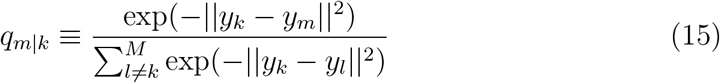
3. SNE then minimizes a cost function, which in this case is the Kullback-Leibler divergence between *p_m|k_* and *q_m|k_* over all data points. To do that, it starts from a random sample of points (Gaussian distributed) and then performs a gradient descent, in which each point *y_k_* is moved around depending on the attraction or repulsion from other data points (see Maaten and Hinton, 2008).

T-SNE makes two main adjustments relative to SNE (the rationale behind these two adjustments is extensively discussed in Maaten and Hinton, 2008). First, it uses a symmetric measure of similarity between two data points, as the joint probability

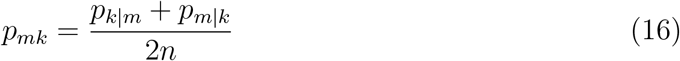

Second, it uses a Student’s t-distribution with one degree of freedom instead of a Gaussian for the low dimensional counterparts.

### 4.4 Measures for cluster evaluation

#### 4.4.1 ARI

The ARI (Adjusted Rand Index, Hubert and Arabie, 1985) is a measure of similarity between two data clusterings *X* (here the ground-truth clusters) and *Y* (here the empirical cluster definitions). We note that the points that are labeled as noise by HDBSCAN are, for the purpose of computing the ARI, assigned to a separate cluster. The computation of the Rand Index considers each pair of observations and determines for each pair of observations (in our case epochs) whether they agree between the two data clusterings. Agreement is defined as either: a) falling in the same cluster in *X* and in the same cluster in *Y*, or b) falling in different clusters in *X* and in different subsets in *Y*. The reason why agreement must be defined over pairs of observations is that the subset partitions do not have to be matched between *X* and *Y*, and that both can contain a different number of clusters. Disagreement is defined as falling in the same subset in *X* but a different subset in *Y*, or falling in different subsets in *X* but the same subset in *Y*. The Rand Index is then defined as the ratio of the number of agreements between the data clusterings (i.e. one epoch being assigned to the same data clusterings) over the total number of agreements and disagreements. The Adjusted Rand Index corrects for a bias in the Rand Index, by subtracting the ratio of number of agreements over disagreements that is expected by chance.

#### 4.4.2 Silhouette

The Silhouette score (Rousseeuw, 1987) is one example measure of cluster quality that does not rely on a ground truth cluster assignment, and is therefore applicable in unsupervised settings. The Silhouette score is computed as follows. For each epoch, we compute the average dissimilarity to all the epoch in the nearest cluster (which could be also the noise cluster), *d_k,nearest_*. We then also compute the average dissimilarity to all the epochs in the same cluster as the epoch, *d_k,same_*. We then compute the Silhouette as

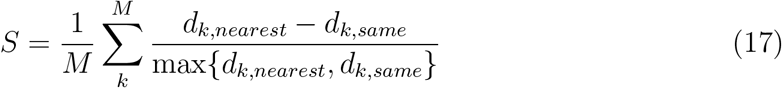

### 4.5 Application to neuronal data

One male macaque monkey performed a passive fixation task while moving bar stimuli (white bars on gray background, 0.25 degrees in visual angle width) were presented. All procedures complied with the German law for the protection of animals and were approved by the regional authority (Regierungspräsidium Darmstadt). Recordings were performed from 64 V1 channels simultaneously, obtained from a chronic Utah array implant (Blackrock). Receptive fields had eccentricities around 3-5 degrees visual angle. We performed band-pass filtering of each channel in the frequency range of action potentials (300-6000Hz) and then thresholded the band-pass filtered signal *x*(*t*) according to (Quiroga, Nadasdy, and Ben-Shaul, 2004), using the threshold 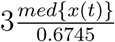, where *med* is the median (i.e. effectively three standard deviations). When the signal *x*(*t*) crossed this threshold, we denoted a spike. After the detection of a threshold crossing, further threshold crossings were suppressed for 0.75ms. Moving bar stimuli were presented in four cardinal directions. Each stimulus bar was presented 20 times. We then pooled all 80 trials together, and added 80 trials containing spontaneous activity. Our aim was then to recover the separate stimulus conditions using unsupervised clustering with SPOTDisClust. We use *n_pts_* = 3 with leaf selection for the HDBSCAN parameters.

## Acknowledgement

Dr. Michael Schmid, Dr. Katharine Shapcott, Dr. Joscha Schmiedt, Dr. Richard Saunders, Dr. Pascal Fries, Cem Uran and Alina Peters were responsible for the collection and preprocessing of the experimental monkey V1 data, with financial support from DFG Emmy Nother grant 2806 (Dr. Michael Schmid). We thank Dr. Felix Effenberger for inspiring discussions on this topic and helpful comments. LG was financially supported by the Erasmus Plus Traineeship Program. FPB was financially supported by the European Union FP7 Project 600925 “Neuroseeker”. MV was financially supported by the Ernst Strüngmann Institute for Neuroscience in Cooperation with Max Planck Society, Frankfurt am Main, Germany.

---

**Figure S1:**
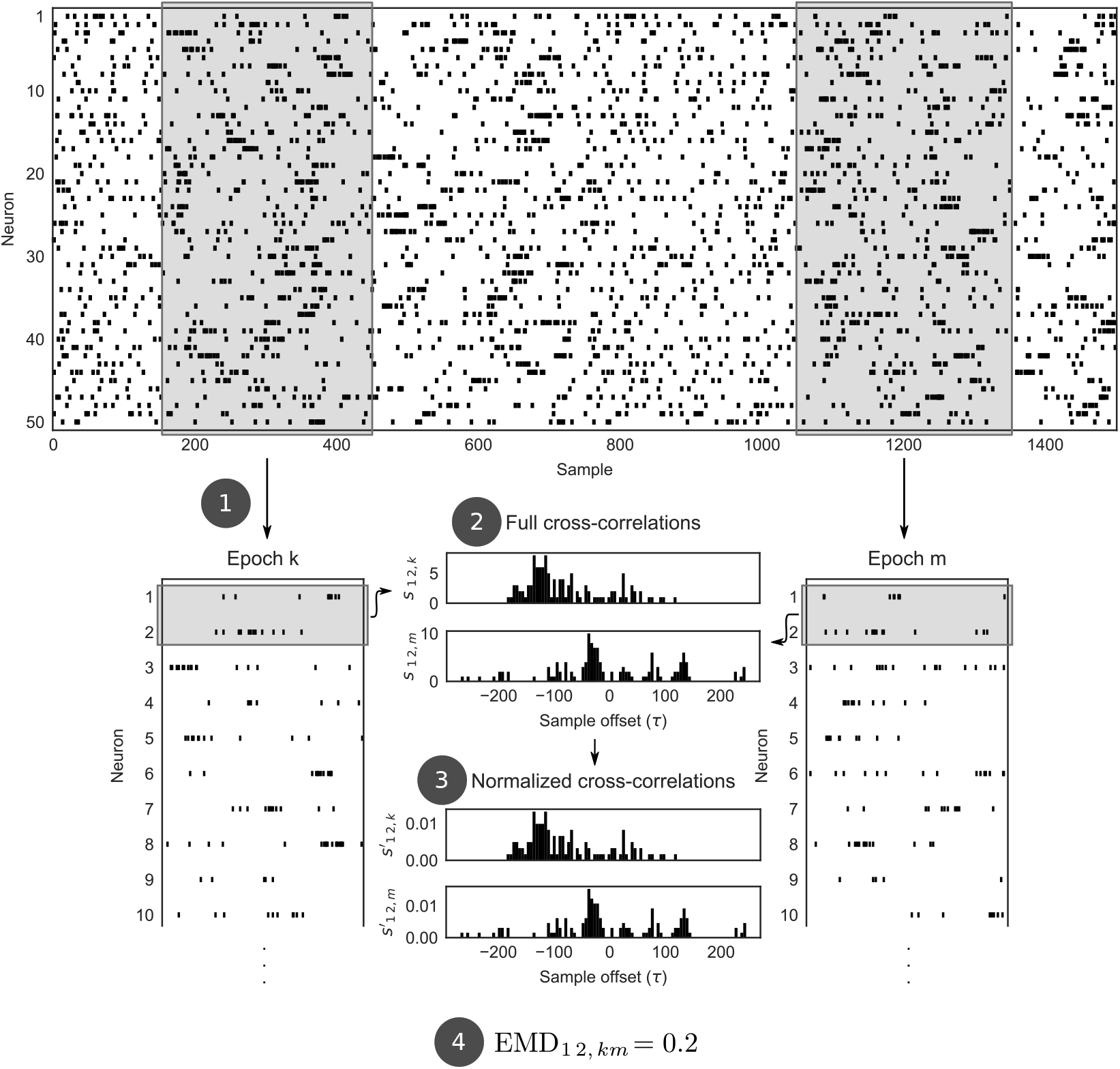
Illustration of the SPOTDis computation. For each epoch (1), the crosscorrelation is computed for each pair of neurons (2). These cross-correlations are then normalized to unit mass (3). For each pair of epochs and pair of neurons (4) we then compute the EMD. The EMDs are then averaged over all eligible neuron pairs (i.e. pairs of neurons active in both epoch *k* and *m*) to compute the SPOTDis between epochs *k* and *m*.

**Figure S2:**
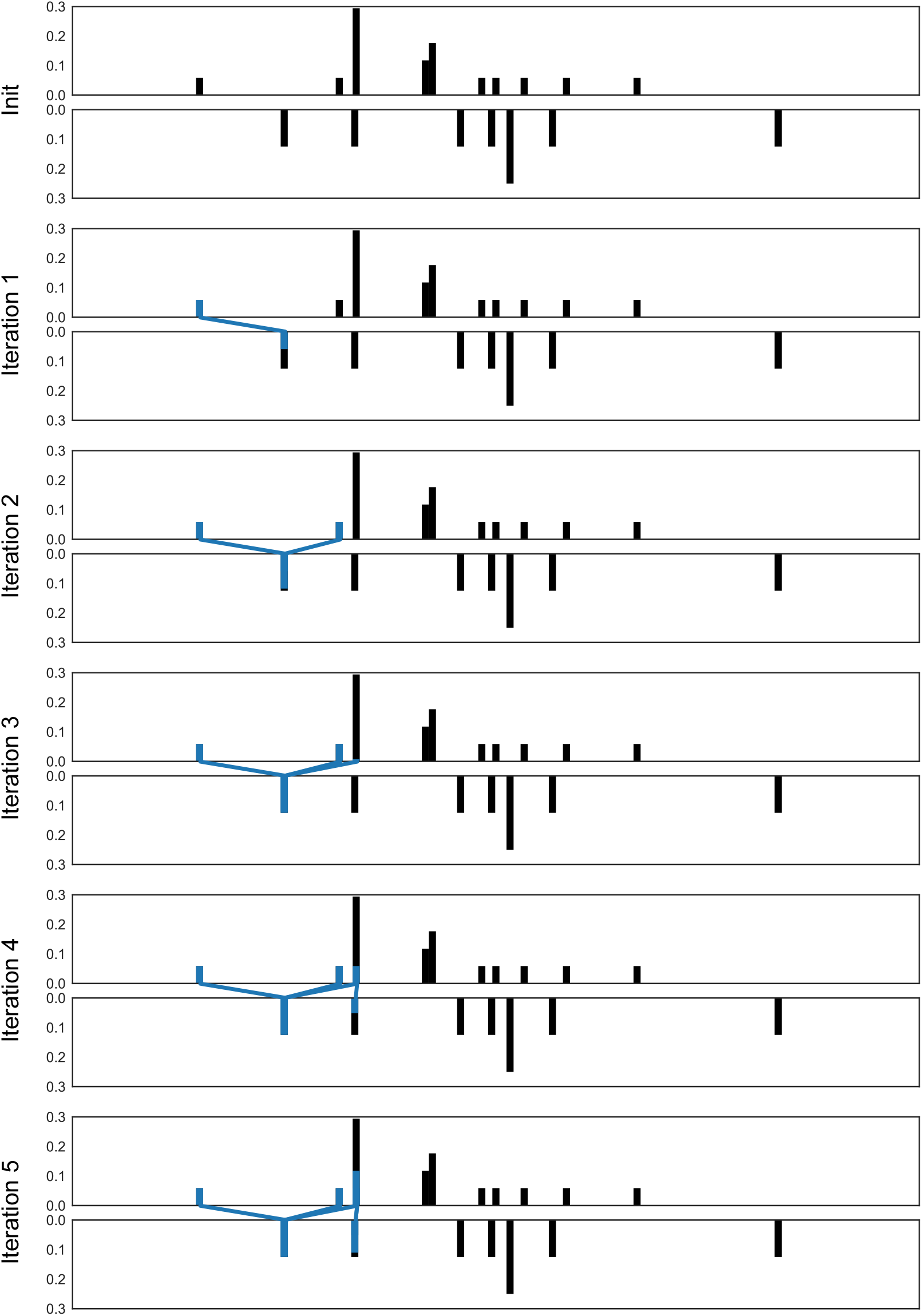
Illustration of the EMD computation. Showing the first 5 iterations of the EMD algorithm on simple example cross correlation histograms. Each normalized crosscorrelation (top and bottom) belongs to one epoch. In each step the mass is transported from the top epoch to the highlighted parts of the histogram and the histogram entries between which mass is transported are connected.

**Figure S3:**
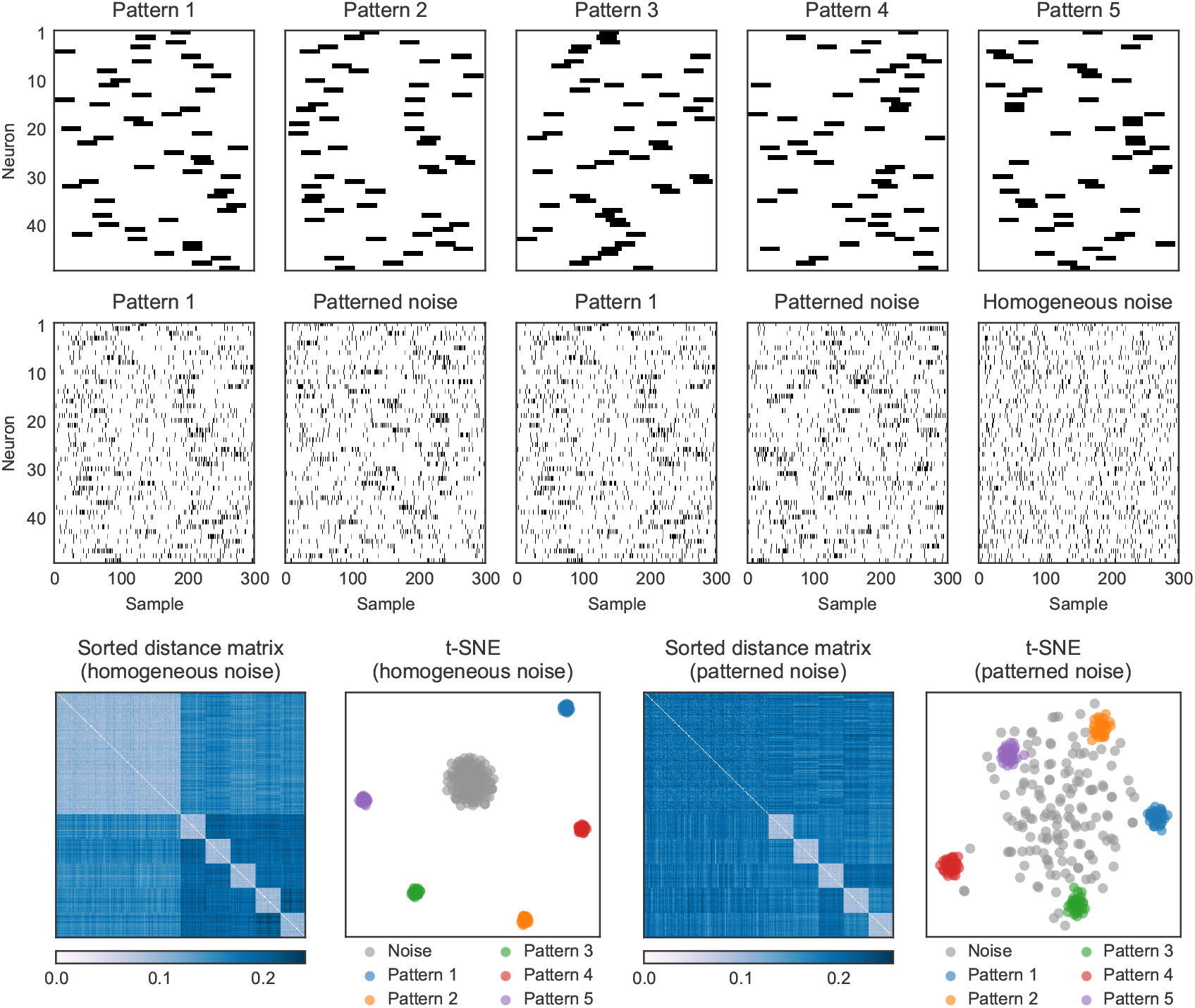
Example figure to illustrate the difference between homogeneous noise and patterned noise. Simulation parameters were *λ_in_* = 0.35 spks/sample, *λ_out_* = 0.05 spks/sample, *T_epoch_* = 300 samples, *T_pulse_* = 30 samples. Homogeneous noise was generated according to a homogeneous Poisson process, while each patterned noise epoch was an instantiation of a unique pattern, that was randomly generated with the same statistics as the four recurring patterns (i.e. for each neuron it had the same values of the pulse duration, *λ_in_* and *λ_out_*). The t-SNE embedding shows that homogeneous noise forms a separate cluster, while patterned noise does not.

**Figure S4:**
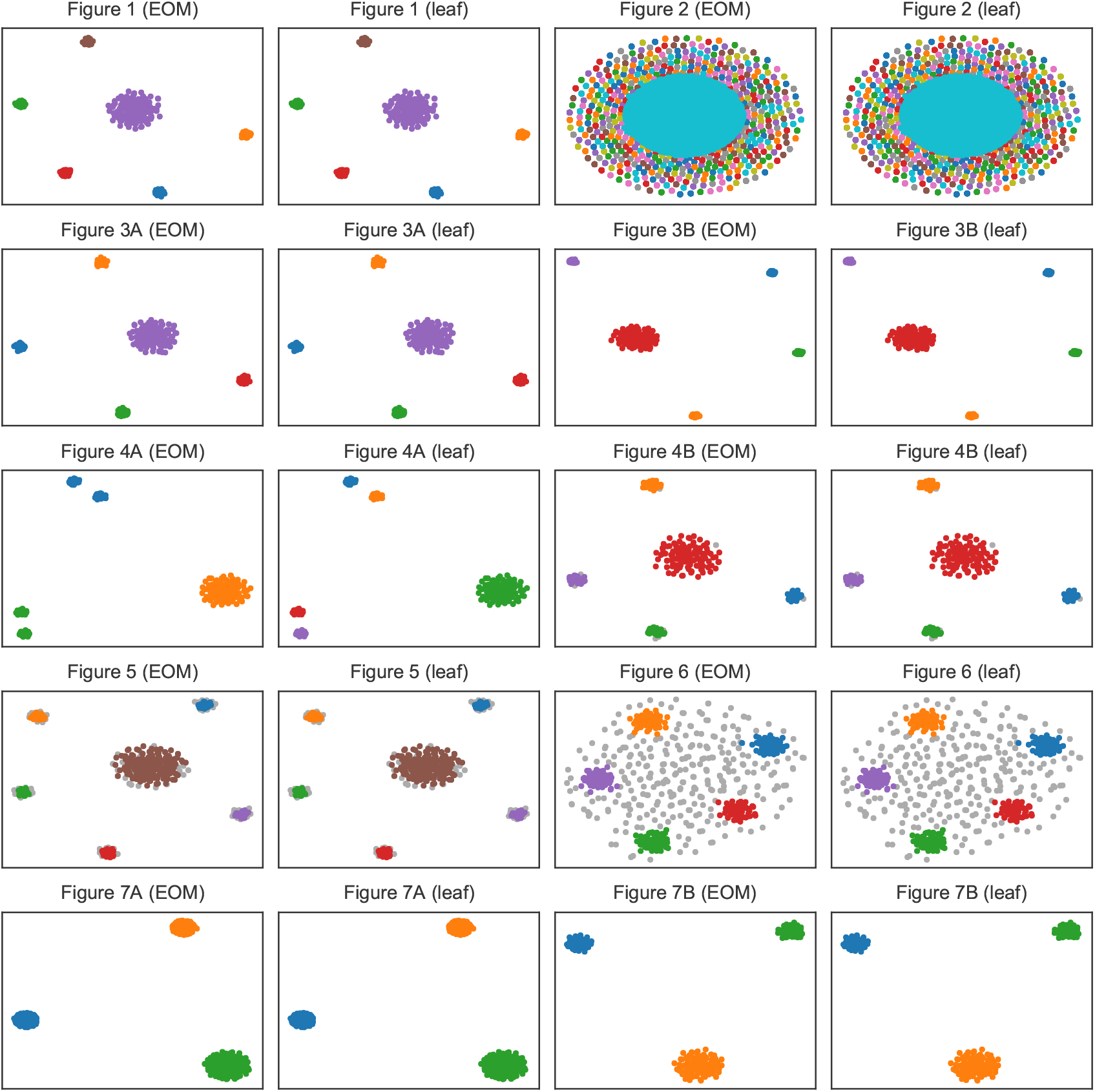
HDBSCAN labels for each of the seven main figures. Shown are the excess of mass (EOM) and the leaf cluster selection methods of the HDBSCAN algorithm (see Methods). Grey points are identified as noise points by HDBSCAN, other colors correspond to clusters.

**Figure S5:**
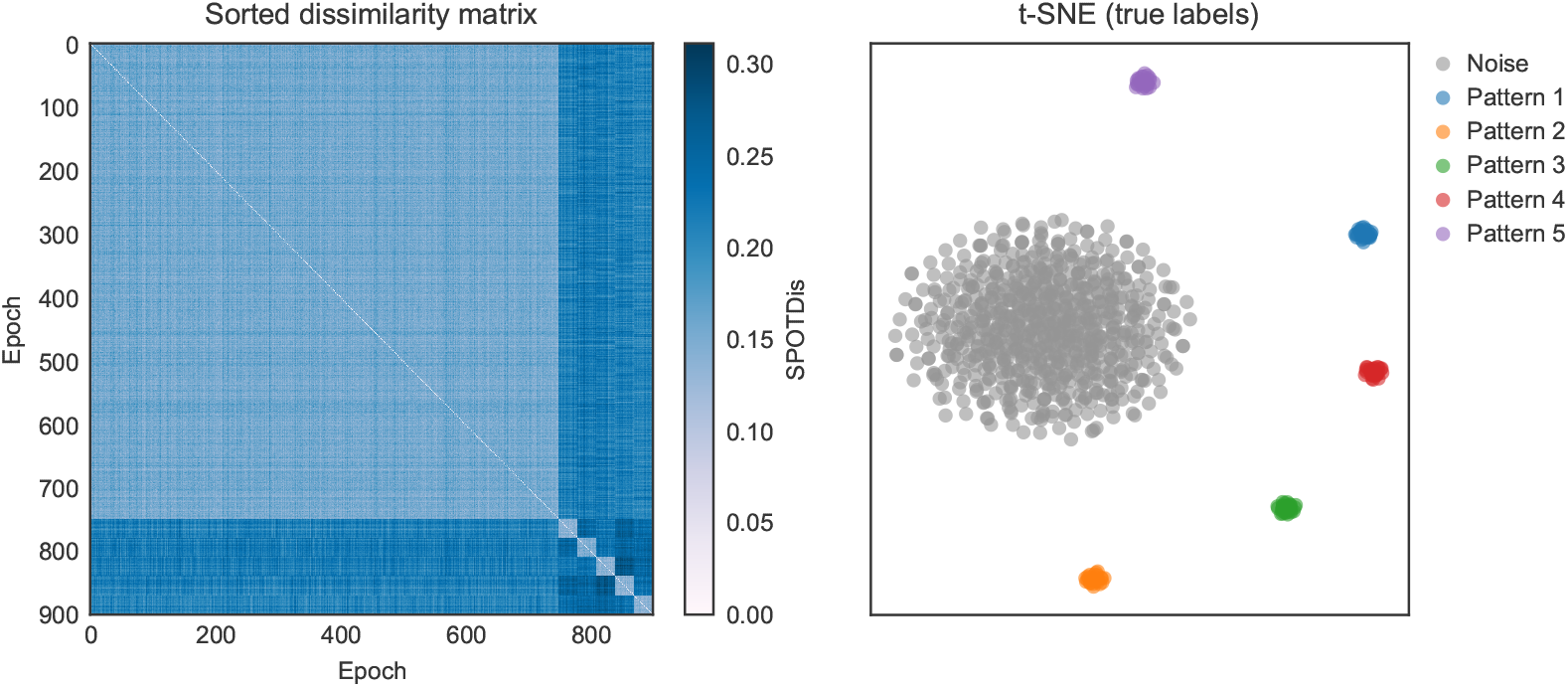
SPOTDisClust performance for cases where there are many noise epochs and few cluster epochs. Same settings as in Figure 1, but now using 750 noise epochs and only 30 epochs per cluster. Even though there is a very large number of noise epochs, t-SNE embedding is still able to reveal the separate clusters.

**Figure S6:**
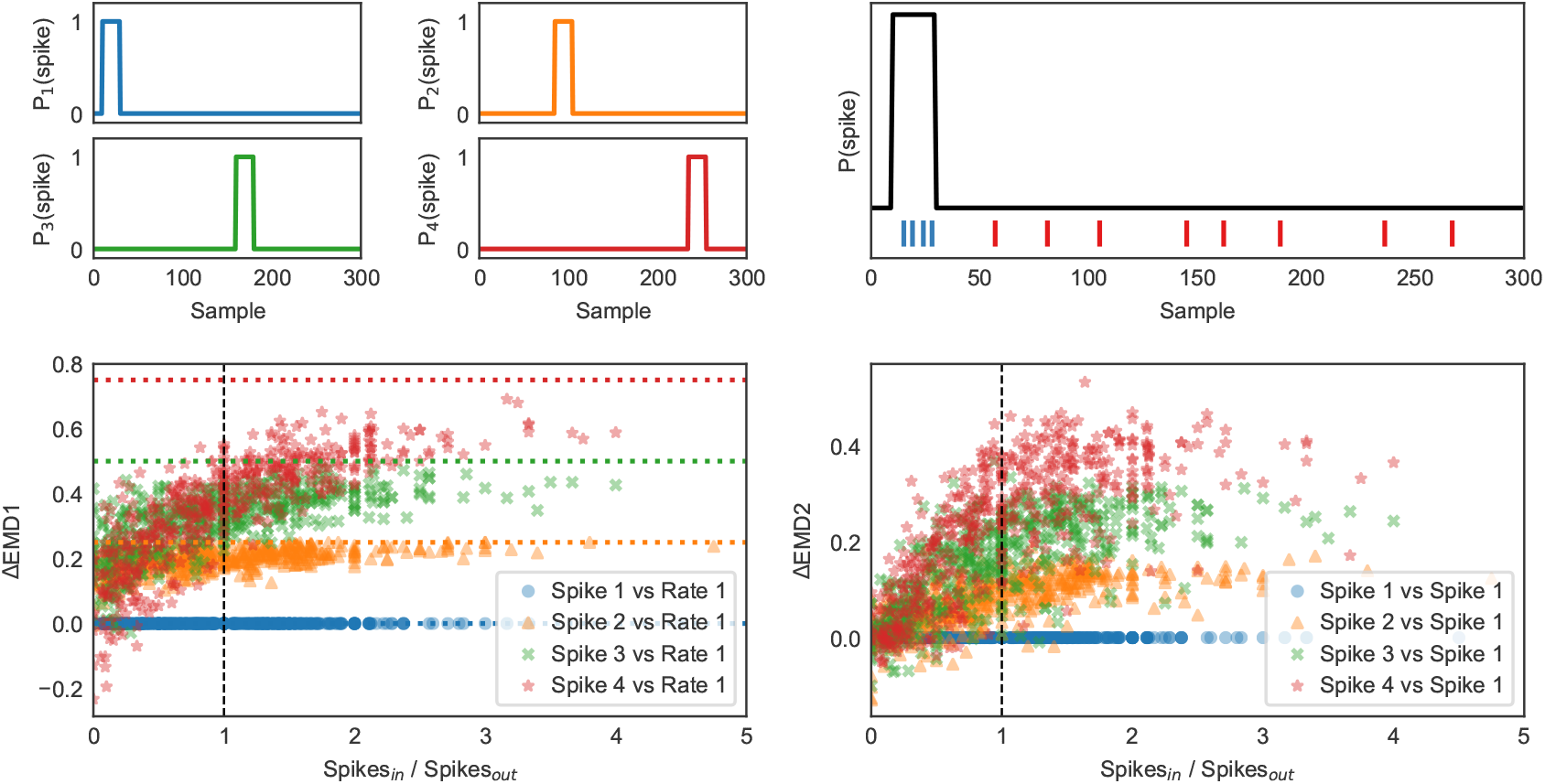
Illustration of how the EMD is a noise-robust measure of distance for spikes. We consider four rate modulations (top left) each consisting of a single activation pulse and a baseline. We then generated spikes from the four rate modulations according to an inhomogeneous Poisson process (top right). The rate modulations may represent crosscorrelations for neuron pairs or activation patterns for single neurons. We then varied the spike rate inside the activation period and outside the activation period, thereby changing the ratio of spikes inside the pulse period to the amount of spikes outside the pulse period (changing the SNR). We then computed the EMD between the normalized mass of rate modulation *k* and the spike realization of rate modulation 1, for all *k* = 1,…, 4, defined as *Δ_k_*. Here the pattern mass can be interpreted as an average of many spike train realizations of that rate modulation. The figure on the bottom left shows, across many spike train realizations of rate modulation 1, ΔEMD(*k*) = Δ_*k*_ – Δ_1_ The dashed lines indicate the EMD between the different patterns. Thus, a value of ΔEMD(*k*) > 0 indicates that the spike train has a lower EMD to its own corresponding rate modulation than to another rate modulation, i.e. it indicates that the assignment of spike train to rate modulation based on the EMD would be correct. We find that as long as the spike count SNR - which we define as the ratio of spikes inside the pulse period to the amount of spikes outside the pulse period - exceeds 1, the minimum value of Δ_k_ occurs when *k* = 1, that is when the EMD is computed between the realization and its corresponding rate modulation (Figure S6). Even when the SNR drops significantly below 1, this holds true. Thus, the proximity of a rate modulation to its realization, in terms of EMD, is highly robust to the insertion of noise spikes around the activation pulse period. We find that the same principle holds approximately true when comparing the EMD between two spike train realizations from the same rate modulation vs. spike train realizations from other rate modulation (bottom right). The robustness to insertion of spikes around the activation period can be intuitively understood from the behavior of the EMD transport distance: When comparing e.g. a realization of rate modulation 1 with the mass of pattern 2, we need to move points out of the pulse period of rate modulation 1, move points into the activation period of rate modulation 2, as well as spreading out spike energy to minimize the distance to the baseline. In contrast, when comparing spikes to the same rate modulation, the transport cost only consists of moving a few points in or out of the pulse period, and spreading the points around to minimize the distance to the baseline.

**Figure S7:**
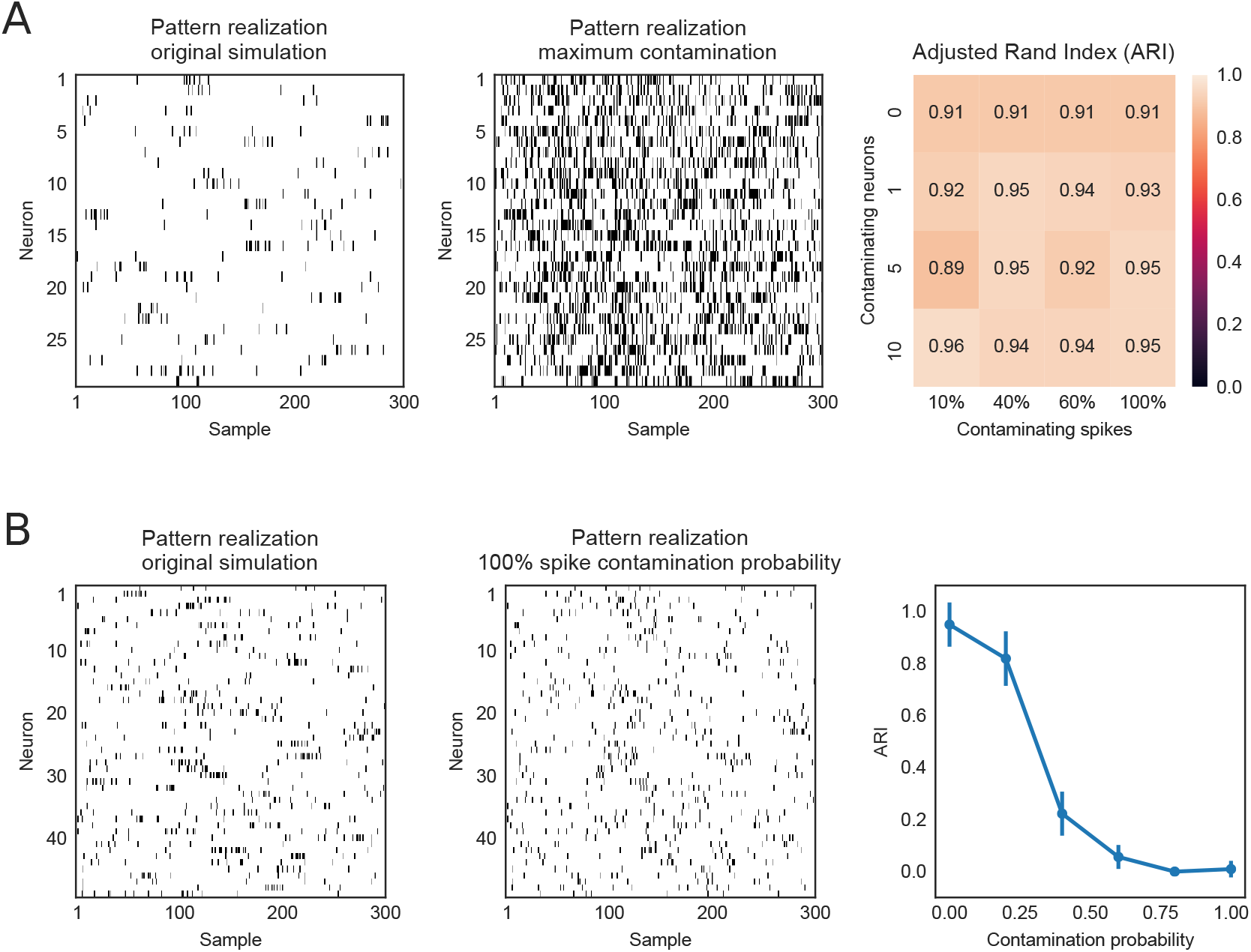
Dependence of clustering performance on spike sorting contamination. (A) Five patterns were generated for a total of 330 neurons, with λ_*in*_ = 0.15 spks/sample, λ_*out*_ = 0.01 spks/sample. Each group of 11 subsequent neurons was assigned to one “virtual electrode”. From this virtual electrode, we then only “recorded” one neuron. The activity of each recorded neuron was then contaminated by randomly inserting spikes from the other “hidden’ neurons. Both the number of contaminating “hidden’ neurons and the fraction of contaminating spikes, as compared to the total number of spikes of each “hidden” neuron, was varied. For example, a value of 5 contaminating neurons and 50% contaminating spikes indicated that 50% of spikes from each of the 5 contaminating “hidden” neurons was added to the recorded neuron. On the left and middle, an original realization of a spike pattern and the observed spike trains after contamination (10 contaminating neurons and each “hidden” neuron contaminating with 100% of its spikes). The table on the right shows the clustering performance compared with the ground-truth (ARI) as a function of spike contamination parameters. B) Patterns were defined using the same firing statistics as in Figure 1. Each group of 5 subsequent neurons was assigned to a “virtual electrode”. The output of all neurons was observed in this case. For each epoch, we then randomly exchanged spikes among the 5 neurons. A contamination probability of 0. 25 meant that each spike had a probability of 0.25 to be transferred to another randomly chosen neuron, i.e. inserted into the spike train of that other neuron. On the left and middle are respectively shown the original spike train output, and the resulting spike train pattern after contamination. On the right is shown the HDBSCAN clustering performance compared to ground-truth (ARI) with standard deviations (across 5 repetitions of the simulation).

**Figure S8:**
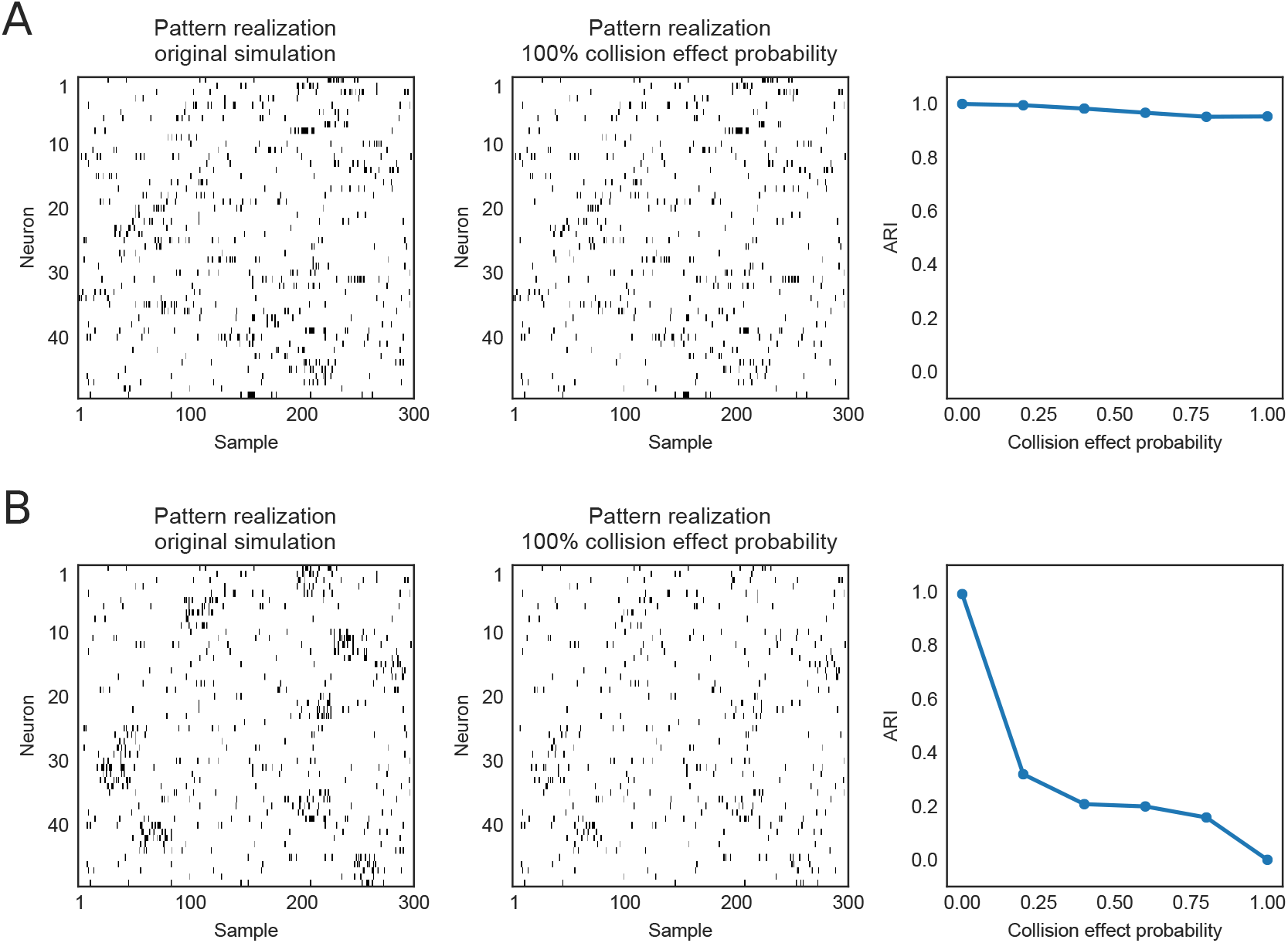
Dependence of clustering performance on spike sorting errors resulting from collisions. The simulations are based on the same firing statistics reported for Figure 1. Neurons were divided into 10 groups of subsequent 5 neurons, each constituting a “virtual electrode”. Whenever two neurons fired simultaneously (i.e. at the same sample) within the group of 5, we removed the two spikes with a certain collision probability. The spike rasters on the left and middle panel show, respectively, an original spike train realization from a pattern and the spike train realization after removing the colliding spikes with 100% collision probability. The panel on the right shows HDBSCAN clustering performance compared to ground-truth (ARI) as a function of collision probability, with standard deviations. Same as (A), but now all neurons within a group of 5 fire according to identical patterns, which strongly increases the amount of collisions, and deteriorates clustering performance as collision probability increases.

**Figure S9:**
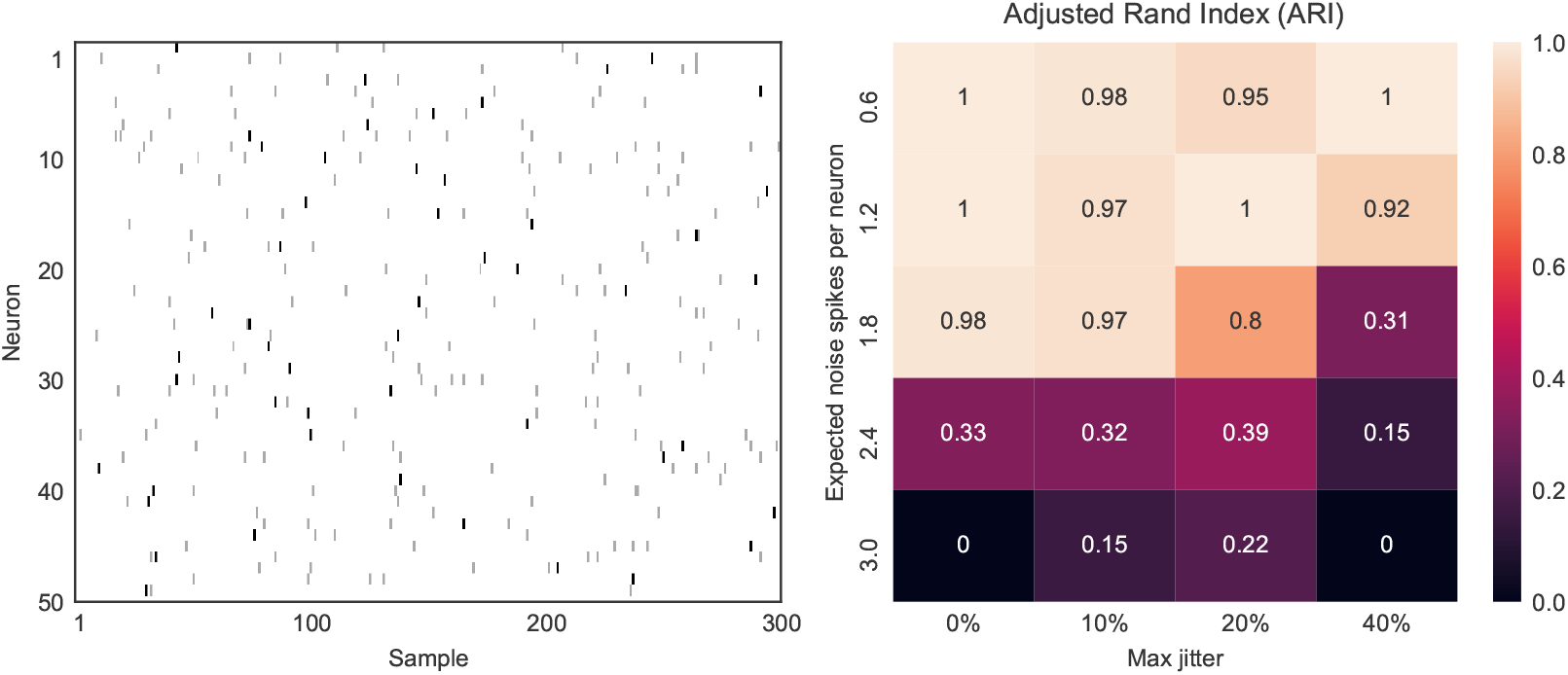
Clustering performance for a case with precise spike sequences. Five patterns were defined by selecting for each pattern a random time point at which the neuron would fire a spike. In addition, we inserted a varying number of noise spikes per neuron according to a homogeneous Poisson process, and also varied the amount of temporal jitter in the pattern spikes. The left panel shows an example for one such pattern, where the pattern spikes are displayed in black and the noise spikes are colored gray. The right panel shows the HDBSCAN cluster quality as compared to ground truth (ARI) with a varying number of noise spikes along with applying an increasing amount of temporal jitter to the pattern spikes. The maximum jitter denoted on the x-axis is defined as a percentage of the total interval length, from which a perturbation value is chosen with uniform probability, for each pattern spike individually. Spikes that are perturbed such that they fall outside of the epoch’s time interval are still considered to be part of the epoch.

**Figure S10:**
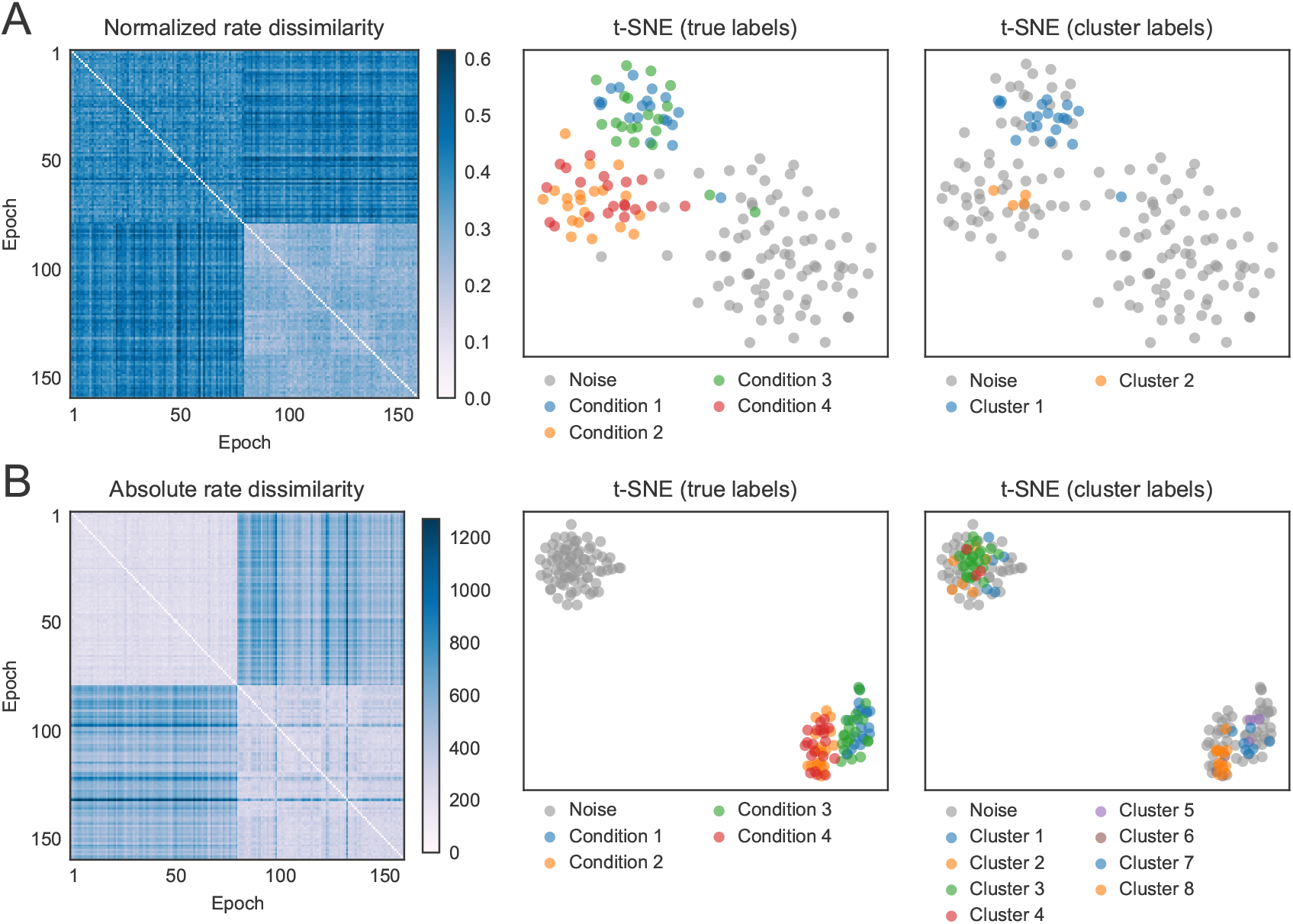
Application to neuronal data, matching Figure 8. For each multi-unit, we computed the spike count in the same temporal window as used for the SPOTDisClust clustering, denoted *r_ik_* (epoch *k*, unit *i*). In (A), we then constructed a normalized population vector as 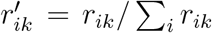 for each multi-unit *i*. We then constructed all pairwise distances between epochs *k* and *M* as the L1-norm among these normalized population vectors, 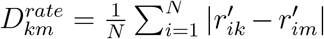. Based on these pairwise distances, we then performed low dimensional t-SNE embedding and HDBSCAN clustering. Shown are the distance matrix, as well as the t-SNE with true labels and cluster labels. The clustering and low-dimensional embedding is unable to separate out all four stimulus directions. In (B), we defined the pairwise distances as the L1-norm on the absolute spike counts, i.e. 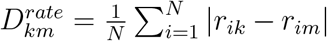, and then performed t-SNE and HDBSCAN clustering. Like in (A), this clustering procedure fails to separate all four stimulus directions.

**Figure S11:**
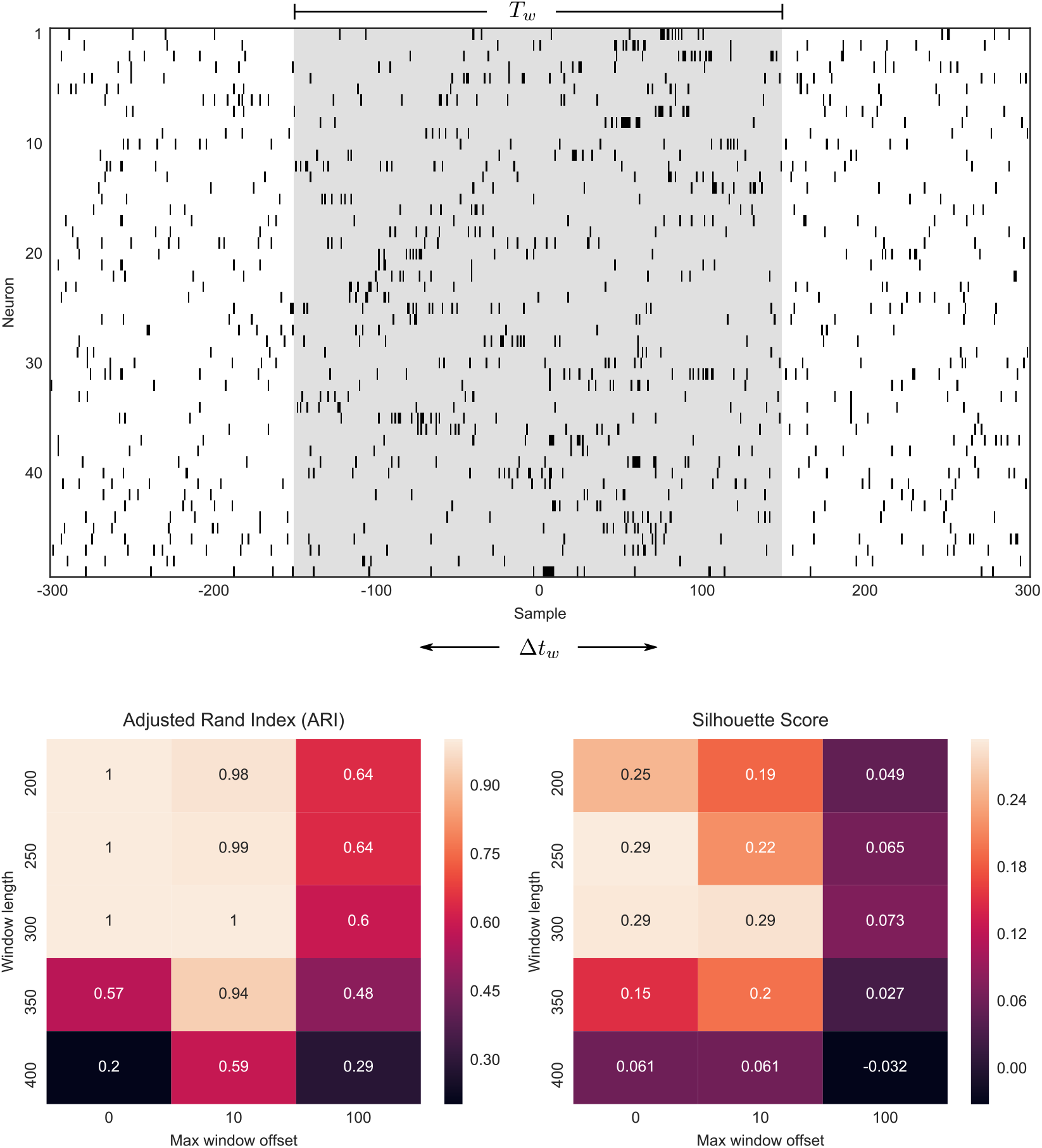
Dependence of clustering performance on chosen window length and temporal jitter of spike pattern onset. The parameter settings for the patterns underlying this figure are equivalent to Figure 1. Each pattern has a length of 300 samples, and is embedded in a larger window starting from −300 samples to +300 samples, with homogeneous noise surrounding the pattern on the left and right. The onset of the pattern is −150 samples plus some random offset Δ*t_w_*. For each epoch realization, the value of Δ*t_w_* was randomly chosen with uniform probability from an interval determined by the maximum window offset (max offset of 100 meant that Δ*t_w_* ∈ [–100,100]). A value of Δ*t_w_* = –50 then meant that the sequence started at −200 samples and ended at 100 samples. For the clustering, we then assume that the sequence duration is unknown. We select a window ranging from –*T_w_*/2 to +*T_w_*/2 samples of length *T_w_*. In case of *T_w_* = 300 and no offset (*Δt_w_* = 0), the generated data matches the simulation of 1. Clustering performance was measured relative to ground-truth (ARI) and with an unsupervised performance measure, Silhouette (see Methods). Clustering performance decreased as the maximum window offset increased, due to the inclusion of noise spikes around the spike pattern. Clustering performance peaked around *T_w_* = 300 samples for both ARI and Silhouette as *T_w_*. indicating that we can determine the “optimal” window length in an unsupervised manner when the sequence duration is unknown.

**Figure S12:**
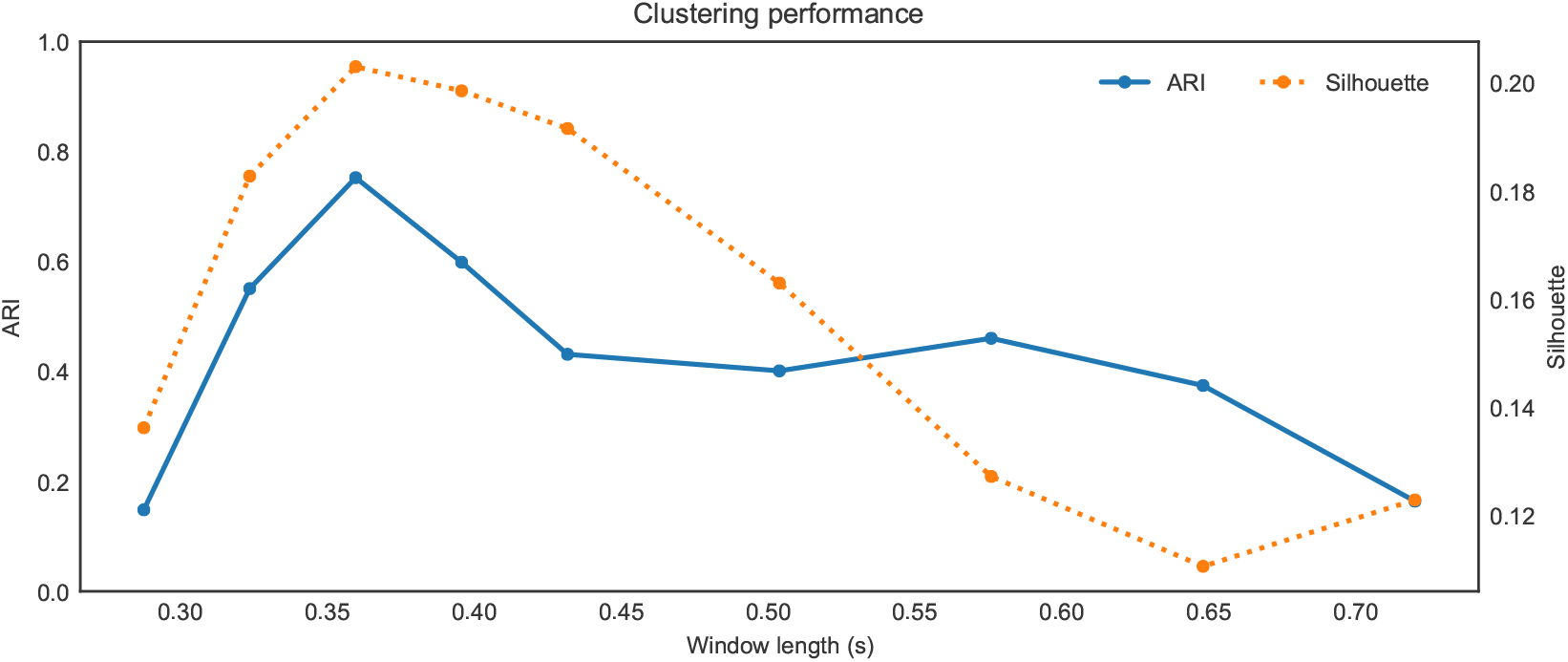
Application to neuronal data of window length selection, matching Figure 8. We repeated the clustering with different window length selections. Windows were centered on 180 ms, the center of the 360 ms stimulation period. Clustering performance, measured both with a ground-truth performance measure (ARI) and an unsupervised cluster quality score (Silhouette), peaked at the window length of 360 ms. Large window lengths lead to an inclusion of baseline firing, decreasing clustering performance, whereas smaller window lengths also deteriorated performance. ARI and Silhouette scores showed a tight correspondence, showing the feasibility of using Silhouette score to “optimize” the window length for sequence detection.

